# ΦX174 bacteriophage viability predicted by protein biophysical modeling

**DOI:** 10.64898/2025.12.01.691700

**Authors:** James T. Van Leuven, Jagdish Suresh Patel, Casey Beard, F. Marty Ytreberg, LuAnn Scott, Kellee Burns, Emma Altman, Yesol Sapozhnikov, Chénangnon Frédéric Tovissodé, Jordan Yang, Holly A. Wichman, Brenda M. Rubenstein, Craig R. Miller

**Affiliations:** Department of Animal Veterinary and Food Science, University of Idaho, Moscow, 83844, USA; Institute for Modeling Collaboration and Innovation, University of Idaho, Moscow, 83844, USA; Department of Chemical and Biological Engineering, University of Idaho, Moscow, 83844, USA; Department of Physics, University of Idaho, Moscow, 83844, USA; Department of Biological Sciences Department, University of Idaho, Moscow, 83844, USA; Department of Chemistry, Brown University, Providence, 02912, USA; Data Science Institute, Brown University, Providence, 02912, USA

**Keywords:** PhiX174, virus, bacteriophage, viability, molecular modeling, amino acid substitutions

## Abstract

The relationship between genotype and phenotype underlies our ability to understand and predict evolution. Efforts to build genotype-phenotype (GP) maps have revealed several unifying rules: epistasis is pervasive, fitness effects are not normally distributed, and the GP map is non-linear and complicated for high-level phenotypes. A critical step in developing GP maps is evaluating how well predictive models do in explaining observed phenotypes. We utilize the simplicity of a bacteriophage (ΦX174) study system to test if an intermediate phenotype (predicted stability of the G capsid protein) explains more complex phenotypes. In doing so, we compare the predictive performance of free energies of folding and binding obtained using numerous molecular modeling methods as well as phylogenetic and basic biochemical/biophysical properties of amino acid substitutions. By creating a large mutational library, we find that ΦX174 tolerates about 50% of the amino acid substitutions we inserted into the G protein and that molecular modeling compliments other substitution models for predicting viability. Mutations predicted to have large destabilizing effects are especially informative and are almost universally detrimental. These large-effect substitutions often coincide with the most conserved residues in the G protein. Apart from large-effect mutations, our ability to predict ΦX174 phenotypes is fairly poor and we explore various potential confounding factors (e.g., codon bias) that could be considered to improve viability predictions.

## Introduction

Understanding how mutations affect organisms is a long-standing goal of evolutionary biology (Alberch 1991). The relationship between genotype and phenotype (GP) determines our ability to calculate substitution probabilities for phylogenetic inference, predict what mutations may occur in the future, and guide how to engineer new proteins. Thus, many methods have been developed to estimate the exchangeability of amino acids in proteins. These methods vary greatly in their sophistication and accuracy. Substitution matrices [summarized in (Kawashima and Kanehisa 2000)] provide a numerical estimate of the substitution probability calculated from transition frequencies in protein alignments. Other simple means of estimating mutational effects use chemical (charge and polarity) and biophysical (size) properties of amino acids to predict a protein’s tolerance for substitutions (Grantham 1974; Guo et al. 2004). More complex models use a combination of approaches to estimate amino acid exchangeability (Bloom et al. 2005; Kumar et al. 2009; Adzhubei et al. 2010). These models may take into account properties of the entire protein (protein stability, location of mutations, solubility, etc.), cellular models of gene expression (Jack and Wilke 2019; Hill et al. 2024), mutational spectra of individual genes (Bloom 2014), and population genetic parameters (Beaulieu et al. 2019) and can offer improved estimates on the effects that mutations have on proteins.

Another method to predict the impact of amino acid substitutions on proteins is to calculate how substitutions change a protein’s folding free energy. The free energy difference between folded and unfolded proteins (ΔG) is a quantitative trait that can be measured empirically from purified proteins or estimated using molecular modeling. Techniques for molecular modeling (e.g., Rosetta, AlphaFold, etc.) have drastically improved, making it possible to predict the impact of large numbers of mutations computationally (Koehler Leman et al. 2020; Pancotti et al. 2022). The difference in stability between wildtype and mutated proteins (ΔΔG) can then be calculated and related to fitness using different statistical approaches. Over the past couple of decades, much effort has gone into testing protein stability as a simple and universal trait for genotype-phenotype mapping (GPM). Several apparent rules have arisen from this work. First, most wildtype proteins occupy a thermodynamically stable position in the local genotype landscape (Bloom et al. 2005; Faure et al. 2024). Many mutations have relatively little impact on protein stability (small ΔΔGs) and very few mutations increase stability of the wildtype protein (Tokuriki et al. 2007). Second, the relationship between fitness and protein stability is non-linear with fitness, decreasing precipitously at particular ΔΔG values. This nonlinearity is derived from thermodynamic theory (Bloom et al. 2005; Wylie and Shakhnovich 2011) and is supported by empirical data (DePristo et al. 2005). Lastly, most proteins unfold when mutations reduce stability by 2-5 kcal/mol (Duan et al. 2020; Li et al. 2020). Thus, protein stability appears to have significant value as a predictor of organismal phenotypes (such as fitness).

However, even with sophisticated computational models, GPM is notoriously difficult. The regulatory networks and protein chaperone systems of cells can mask the effects of amino acid substitutions at the organism level (Fares et al. 2002; Tokuriki and Tawfik 2009), epistatic effects that deviate far from additive make predictions difficult (Weinreich et al. 2005; Sarkisyan et al. 2016; Gonzalez and Ostermeier 2019; Barnes et al. 2022), and high-throughput experimental measures of phenotypes are still challenging to acquire for many organisms. Recent methodological improvements (e.g., deep mutational scanning, CRISPR, etc.) are rapidly improving our ability to measure the phenotypes of many variants (Levy et al. 2015; Venkataram et al. 2016; Li et al. 2020; Huss et al. 2021) but there is still a need to perform GPM on different protein types (structural, enzymes, etc.) as past studies have shown categorical differences in how genotype relates to phenotype (Høie et al. 2022). To address this, we chose a study system with intermediate complexity—where the relationship between protein folding and high-level phenotypes like fitness should be relative direct (Bull et al. 2011; Wylie and Shakhnovich 2011).

ΦX174 is a virulent bacteriophage of *Escherichia coli*. Infectious virus particles are composed of four types of proteins (F, G, H, and J) and a single-stranded circular DNA genome that is 5.3 kilobases in length. Host infection occurs when the spike (G) protein pentamer interacts with the lipopolysaccharide (LPS) layer of potential host bacteria. This interaction causes the dissociation of one G pentamer from the virion, leaving the underlying capsid (F) proteins to bind to the host LPS. This association leads to conformational changes in capsid proteins that cause the pilot protein (H) to extend through the bacterial membrane into the cytoplasm for the translocation of phage ssDNA (Sun et al. 2017). Mutations in these genes can affect viral growth rate and determine which host strains the phage can infect (Crill et al. 2000; Pepin and Wichman 2007; Vale et al. 2012). However, the role of protein stability in determining these phenotypic effects is unknown.

To understand ΦX174’s ability to tolerate mutations and test our hypothesis that protein stability is a useful predictor of fitness, we performed extensive molecular modeling of the ΦX174 spike protein and empirically tested the viability of 420 amino acid variants of ΦX174. We found that nearly half of these variants produced viable phage and that biophysical modeling correctly predicts inviability for mutations having large destabilizing effects. We developed and compared a logistic regression model relating viability to protein biophysics to several models that utilize basic biochemical properties and phylogenetic conservation. We found that sequence conservation information is very informative in predicting viability. However, in instances where taxon sampling is poor (few related sequences available), molecular modeling provides a good approach to estimating the viability of protein substitutions. The utility of molecular modeling is discussed in relation to simpler methods for estimating the effects of amino acid substitutions, such as differences in biochemical properties of amino acids, substitution matrices, and phylogenetic conservation. This study extends our previous experiment that investigated a much smaller set of mutations in a G4-like phage (Lee et al. 2011).

## Results

### Surveying the genotype space of the ΦX174 spike protein

We mutated 21 residues in gene G to all 64 possible codons and sequenced isolates to identify mutations resulting in viable phages. These 21 sites were choosen to include amino acids with different chemical properties, with variable locations in the G proteins (e.g., buried, surface), and with variable conservation among microvirids. Using site-directed mutagenesis with degenerate primers, we created pools of variants for each mutated residue and recovered viable phages by picking plaques. Using the initial starting codon frequencies and a multinomial Bayesian framework (see Methods for details), we determined the necessary sampling depth (number of plaques) to achieve a 0.99 probability of observing all viable mutations at these 21 sites (**Supplementary figure S1**). This resulted in an average of 136 plaques sequenced per mutated site. However, we found 30 mutations with a marginal probability (0.99-0.95) of being observed if viable considering our initial sampling level. These mutants were not observed in our initial screening, so we attempted (in triplicate) to make these mutations using site-directed mutagenesis. 15 of the 30 yielded viable phage. Combining all data (initial screen and site-directed efforts), we observed 223 of the possible 420 (53%) amino acid variants and 614 of the 1344 (46%) possible codon variants (**Table 1, Supplementary data**). The tolerability to mutation varied drastically among the 21 sites. Residues G119 and D45 tolerated all possible amino acids (**Figure 1**). Several other sites tolerated most amino acid substitutions. Residues S74 and D125 could only be mutated to one other amino acid. We then explored properties of these mutations and their utility in predicting viability.

**Figure 1.**
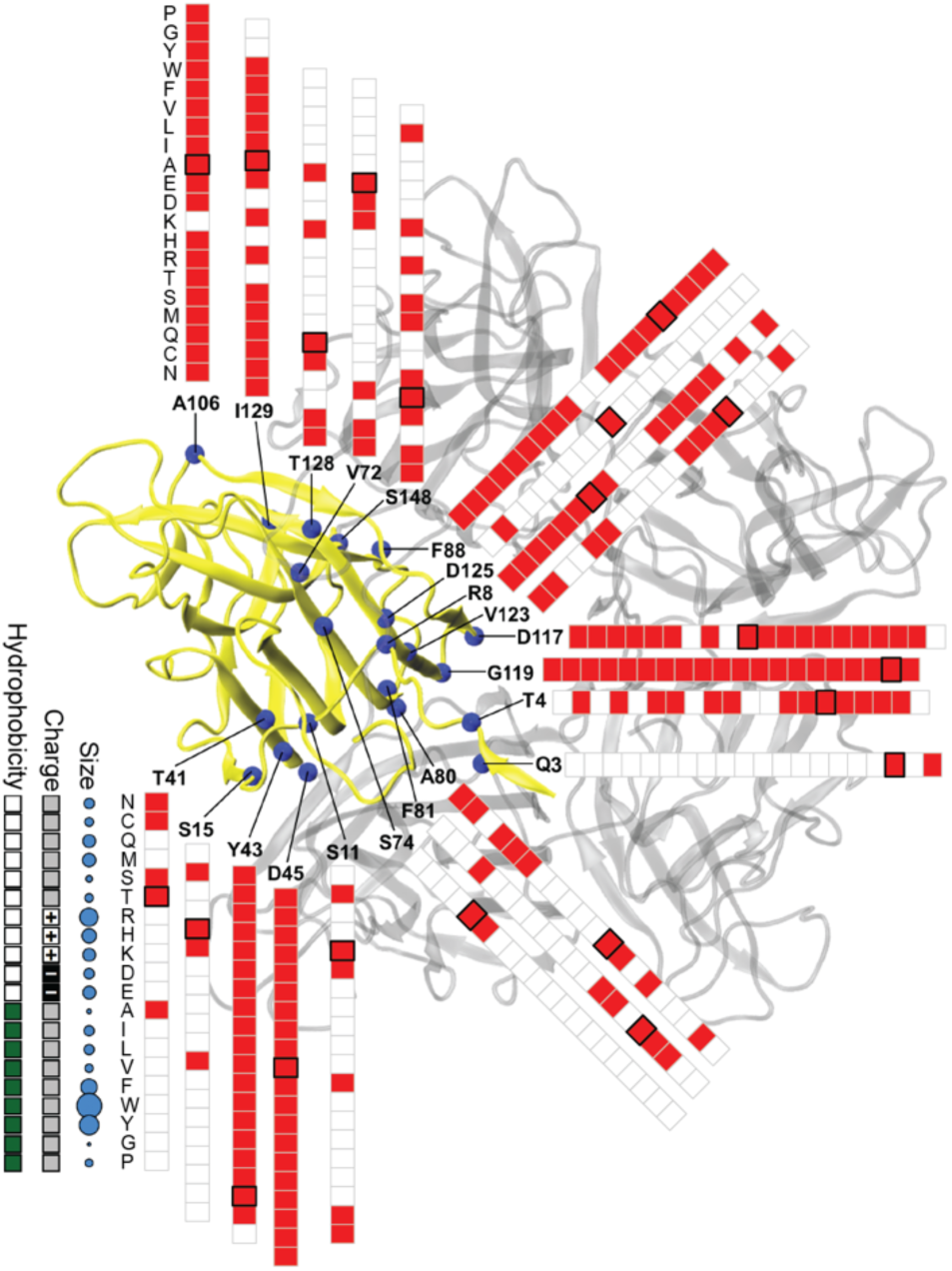
Substitution tolerability is highly variable among mutated residues. The crystal structure of one G protein monomer is show in yellow with the pentamer structure in grey. Viable (red), inviable (white), and wildtype (black box) substitutions are shown along with general amino acid properties.

**Table 1.** List of residues mutated in this study. The number of plaques from each residue that were sequenced; #Plaques, the number of viable codons recovered; #Cods, and the number of viable amino acids recovered; #AAs is shown for each residue.

| Residue | #Plaques | #Cods | #AAs |
| --- | --- | --- | --- |
| Q3 | 91 | 4 | 2 |
| T4 | 247 | 38 | 13 |
| R8 | 177 | 44 | 15 |
| S11 | 126 | 21 | 6 |
| S15 | 160 | 15 | 4 |
| T41 | 107 | 14 | 5 |
| Y43 | 166 | 54 | 19 |
| D45 | 165 | 47 | 20 |
| V72 | 111 | 12 | 6 |
| S74 | 160 | 10 | 2 |
| A80 | 122 | 27 | 9 |
| F81 | 89 | 10 | 6 |
| F88 | 203 | 38 | 14 |
| A106 | 138 | 50 | 19 |
| D117 | 159 | 43 | 17 |
| G119 | 204 | 45 | 20 |
| V123 | 161 | 33 | 9 |
| D125 | 172 | 4 | 2 |
| T128 | 122 | 22 | 9 |
| I129 | 124 | 28 | 15 |
| S148 | 107 | 26 | 10 |

### Defining viable versus inviable

We classify phages as inviable if they are unable to produce infectious phage particles that we can detect. Variants that are extremely inefficient at forming plaques and/or form very small plaques may be viable but evade our detection limits. Thus, we took several steps to have high confidence that what we call inviable variants are indeed extremely unfit. We sequenced the DNA variant library and compared variant frequencies to observation frequencies of recovered isolates. We found that, while most variants are observed in proportion to their expected ligation-mix frequency (p≤0.01, r=0.91), there are nine significant departures from expectation (seven below and two above, **Supplementary figure S2-3**). These seven variants demonstrate that low plating efficiency variants probably exist but at surprisingly low frequency in our “viable” mutants. However, because the initial variant library is pooled (proportional data) we cannot calculate absolute plating efficiency values but can calculate plating efficiency relative to the wildtype. Given the high correlation between expected and observed frequencies, we suspect that the plating efficiencies of most observed variants are close to that of the wildtype. We also measured the plaque size for the eight infrequently observed variants and found they produce small plaques (**Supplementary data**, t-test, p < 0.05). Among all codon variants that result in these eight amino acids, 73% (11/15) form plaques smaller than those formed by the wildtype compared to 40% (156/389) among all other variants (p = 0.02, test for equal proportions, see Methods). Similarly, 80% (24/30 of the codons) of the plaques from the 15 amino acid variants that we made specifically using site-directed mutagenesis were smaller than those formed by the wildtype—a significantly greater proportion than the other variants (p=4.8E-5, test for equal proportions, see Methods).

### Protein modeling with FoldX and PyRosetta

To investigate the relationship between protein folding stability and viability, we used FoldX and PyRosetta to predict the thermodynamic effects of all single amino acid substitutions on the folding and binding stability of protein G of ΦX174. This was done by calculating the difference in Gibbs free energy (ΔG) between folded and unfolded or bound and unbound variants of protein G (**Figure 2**). ΔΔG_bind_, the difference in ΔG between the mutant and wild type proteins, was not strongly influenced by whether it was calculated from the binding of monomers into a dimer, a monomer and dimer into a trimer, or a tetramer and a monomer into a pentamer (**Supplementary data**). The Pearson’s correlation coefficient (r) between the different arrangements was always high (**Supplementary figure S4**): FoldX dimer to trimer r=0.96, dimer to pentamer r=0.97, trimer to pentamer r=1.0; PyRosetta: dimer to trimer r=0.88. This strong coorelation could be a result of FoldX and PyRosetta poorly predicting non-local effects on protein structure. FoldX and PyRosetta are not well correlated with one another: PyRosetta dimer to FoldX dimer r=0.31, PyRosetta trimer to FoldX trimer r=0.31, PyRosetta fold to FoldX fold r=0.40. There is, however, high variability in the correlation between PyRosetta and FoldX among different residues (**Supplementary figures S5 and S6**). For example, the correlation coefficient (r) between FoldX dimer binding and PyRosetta dimer binding for residue S15 is 0.92, while for residue D117 it is 0.015. Thus, for some residues FoldX and PyRosetta make very similar predictions but for others the predicted effects are completely different. The seemingly low correlation overall between PyRosetta and FoldX has been noted in past studies and is attributed to the methods, differing abilities to relax the protein structure and allow for atoms with different volumes (Yang et al. 2020).

**Figure 2.**
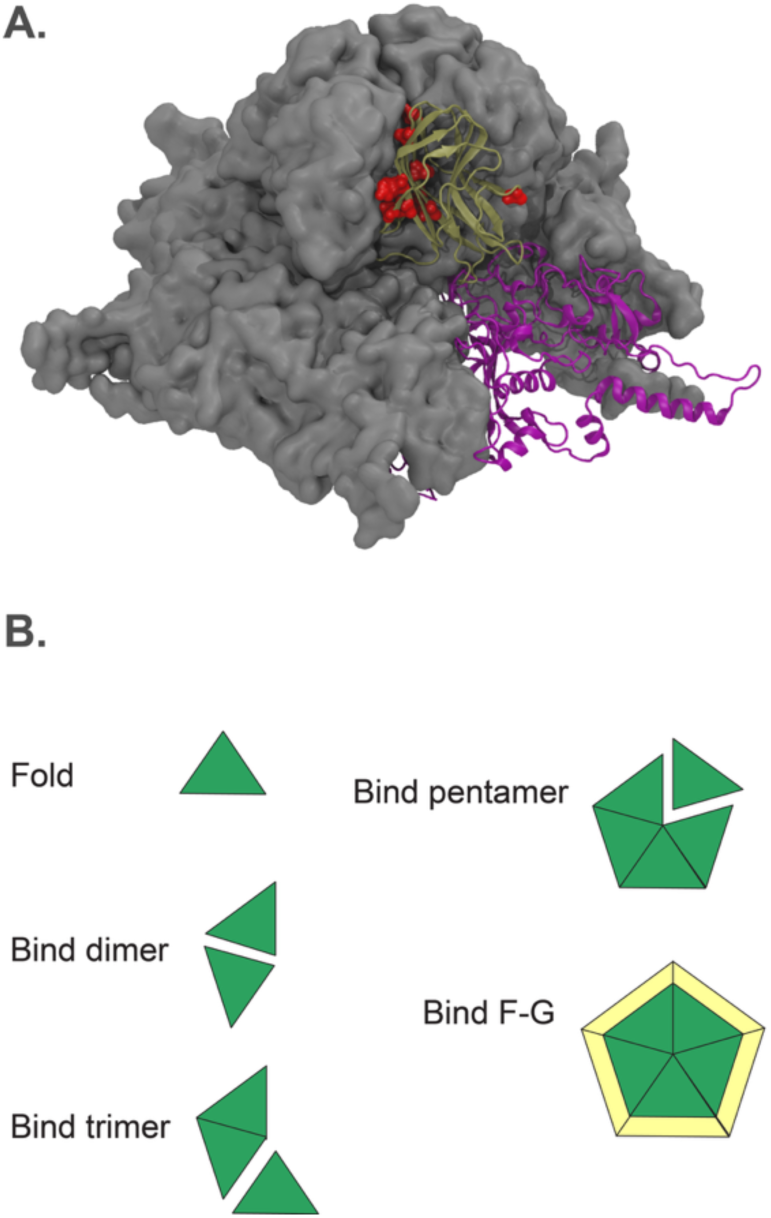
Structural model of ΦX174 G (gold) and F (purple) proteins (A). One monomer of each is displayed in ribbon format. Other proteins in the pentamers are shown using a space-filling display. Mutated residues are shown in red. (B) illustrates the five different ΔΔG quantities measured. These include folding stability and four ways to model protein binding. For example, for “Bind pentamer” the ΔΔG effects were calculated for a protein structure that included all monomers. For “Bind dimer”, the structure only included two monomers.

### Statistical modeling of protein stability

To compare the viability of ΦX174 variants to their predicted stability effects, we developed a multistage binomial (MSB) model, an extension of the traditional binomial model such as logistic regression (see Methods for details). Logistic regression is often used to model binary response variables (e.g., viability). Our model differs from the traditional logistic model in a couple of ways. First, the logistic curve is allowed to asymptote at values < 1, thus assuming that not all mutations with negligible effects on protein stability will be viable. Second, we allow predictors to interact multiplicatively (instead of additively) such that if either folding or binding stability are sufficiently destabilized, the effect in combination is a low probability of being viable. The model also accounts for predictor (ΔΔG) errors, which we recently showed to be ±2.9 kcal/mol for FoldX folding and ±3.5 kcal/mol FoldX binding estimates (Sapozhnikov et al. 2023).

Using the MSB model, we fit ΔΔG values to empirical measures of ΦX174 viability and compared models with the following predictors (alone and in select pairwise combinations): PyRosetta folding, dimer binding, and trimer binding; FoldX folding, dimer binding, trimer binding, and pentamer binding; and FoldX F-G binding (**Table 2**, **Figure 3**). None of the molecular models predicted viability well. The model with the best fit based on AIC and r^2^ is the bivariate model that uses FoldX trimer binding and FoldX folding (r^2^ = 0.273). During the analysis, we observed that residue 119 is predicted by FoldX to destabilize the folding of the G protein by 4-12 kcal/mol but is extremely tolerable to amino acid substitutions. The effects of substitutions from glycines (like residue 119) to other amino acids are inherently difficult for FoldX to predict because of the molecular clash caused by the side chain of the substituted residue and limited FoldX side chain optimization. The ΔΔG estimates of glycines to other amino acids are typically large (**Supplementary figure S7**). The average van der Waals clash terms of the FoldX predictions for all glycines in the G protein is 6.8 kcal/mol, while the average of other amino acids is 1.2 kcal/mol. All 19 mutations at G119 have clash scores above 2.5 kcal/mol, highlighting likely inaccuracies of FoldX in modeling this residue. PyRosetta’s estimates of ΔΔG for G119 substitutions are much lower than those of FoldX (**Supplementary figure S5**). Excluding G119 from the analysis greatly improves FoldX predictions. PyRosetta always predicts viability poorly, regardless of whether G119 is included (r^2^ = 0.055 for the bivariate model). The remainder of our analysis uses the FoldX trimer binding and FoldX folding models with G119 excluded (results with G119 included are shown in **Supplementary table S1**).

**Figure 3.**
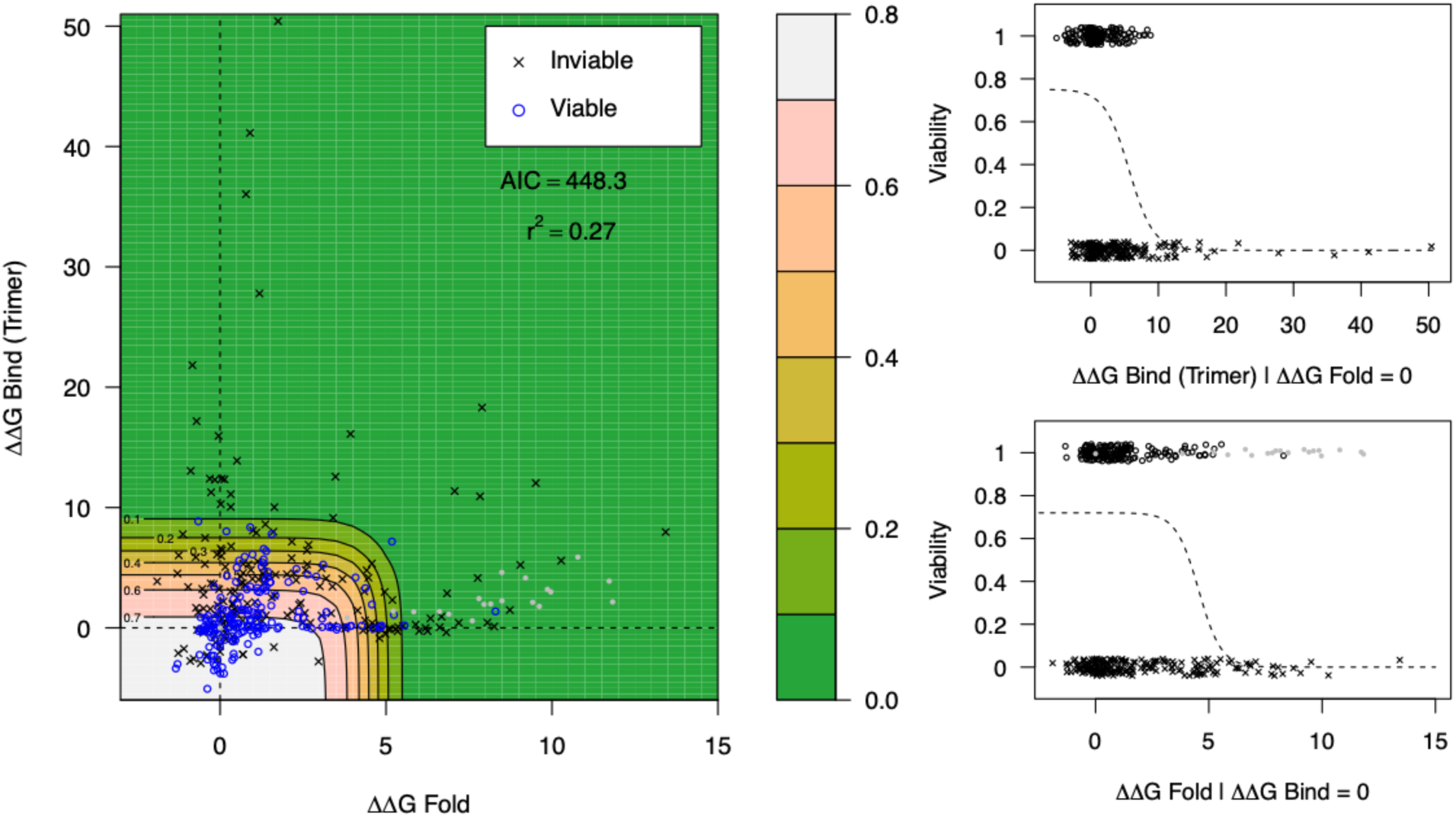
Viability data plotted on top of the best-fit logistic regression model. Variants with large, predicted stability are inviable (x’s), but most mutations have small stability effects. On this small-effect plateau, about 70% of variants are viable. Panels to the right are profiles of the surface conditional on one predictor set to 0–the dashed lines in the heatmap. Gray dots are site 119 variants that were omitted from model fitting.

**Table 2.** Fit of different logistic regression models based on AIC and Nagelkerke’s pseudo-r^2^. Models using FoldX (FX) were run with and without predictor error included (except F-G bind, for which we had no error estimate); PyRosetta (Py) estimates lack error estimates. While inclusion of error is preferable, we provide fits without as well with so fair AIC and r^2^ comparisons between FX and Py can be made. The best model is clearly FX fold + FX bind: trimer.

| Model | AIC with (w/o error) | $r^2$ with error (w/o error) |
| --- | --- | --- |
| FX F fold | 499 (503) | 0.105 (0.094) |
| FX F bind: trimer | 481 (485) | 0.164 (0.153) |
| FX F bind: dimer | 487 (490) | 0.145 (0.136) |
| FX F bind: pentamer | 488 (489) | 0.141 (0.140) |
| FX F-G bind: pentamers | (530) | (0.001) |
| Py fold | (520) | (0.037) |
| Py bind: trimer | (517) | (0.047) |
| Py bind: dimer | (518) | (0.045) |
| FX fold + FX bind: trimer | 449 (459) | 0.273 (0.242) |
| Py fold + Py bind:trimer | (519) | (0.055) |

### Protein instability causes inviability

Despite the low r-values of the MSB model to viability data, there is a clear effect of protein stability on viability, especially for large-effect mutations. Nearly all mutants that are predicted to reduce stability by more than about 5 (10 for trimer binding) kcal/mol are inviable (**Figure 3**). 82% (134/164) of viable viruses have predicted stabilities of less than 2.5 kcal/mol, compared to only 45% (97/216) of inviable viruses. The model illustrates how for both folding and binding, the viability shoulder is in similar locations (right panels in **Figure 3**). If we define the shoulder by where the surface has decreased 25% from its asymptote, this occurs right at ΔΔG = +4 for both folding and binding. The asymptote is around 0.7, which is approximately the proportion of viable to inviable mutants with no predicted ΔΔG effects. We presume that many of these are inviable due to some reason besides changes in protein stability, although, an alternative explanation is that the ΔΔG predictions are commonly incorrect.

### Other predictors of viability

We also investigated how well the viability of ΦX174 mutations can be predicted with information besides protein stability modeling. These properties included change in electrostatic charge, change in molecular weight, change in hydrophobicity, amino acid substitution matrices (PAM30, PAM70, BLOSUM62, and BLOSUM100), and phylogenetic conservation (**Supplementary figure S8**). Of all the methods of predicting viability, those based on amino acid properties require the least amount of prior information of a particular protein because they are calculated from amino acid alignments of large sets of proteins. However, these estimates provide some predictive power on the tolerability of mutation. Mutations with BLOSUM100 score above -2 (high exchangeability) are viable about 70% of the time, which is very similar to ΔΔG modeling asymptote. Mutations with BLOSUM scores below -2 are viable about 50% the time (**Supplementary figure S9**).

Incorporating taxonomic information from other sequenced microvirids helps predict what mutations will be tolerated by ΦX174. We included sequences from the α3, G4, and ΦX174 clades. Among these taxa, gene G is highly conserved. Of the 21 residues that we experimentally mutated in ΦX174, three (Q3, S74, and S148) are completely conserved across microvirids and six residues have only two alleles (**Figure 4**). Among the ΦX-like viruses (subset of 18 taxa), only six residues (8, 43, 45, 106, 117, and 125) have more than one allele. Using the entire microviridae phylogeny, we estimated the conservation of residues in the G protein using rate4site and predicted their mutability using Sorting Intolerant From Tolerant (SIFT) (**Supplementary data**). SIFT predicts that only 73 of the 399 possible mutations will be viable (**Supplementary figure S10**, 0.05 cutoff). Residues Y43 and A106 were predicted to be highly mutable, with 12/19 and 14/19 mutations predicted to be viable. These predictions turned out to be mostly correct. Residues Y43 and A106 are the most mutable sites that we identified by site-directed mutagenesis. However, nearly all mutations (not just 12/19 and 14/19) to these two residues resulted in viable phages. Moreover, residues R8, D117, and G119 were almost equally tolerant of mutations even though SIFT predicted that many of the mutations we made would be inviable. In general, these tools perform better than molecular modeling at predicting the viability of residues with small predicted ΔΔG values (**Figure 5**). At these residues, the small ΔΔG values would indicate all substitutions to be viable, but phylogenetic information provides insight into which specific substitutions are tolerated. Unlike molecular modeling, both tools require sequence alignments and may be susceptible to poor taxon sampling. For example, in our dataset we observed little variation at some residues (e.g., R8, I129) and rate4site and SIFT understandably predicted that few mutations would be viable at these residues. FoldX and PyRosetta on the other hand predict relatively small ΔΔG values for these mutations and thus provide better estimates on their tolerance to substitutions. We used our logistical regression model to compare the performance of models based on amino acid properties, biophysical modeling, and phylogenetic conservation and found that that rate4site outperforms the other metrics for our dataset (**Table 2**). We did not test models that combine biophysical modeling and evolutionary (phylogenetic) models because our primary goal was to determine how well a priori models perform.

**Figure 4.**
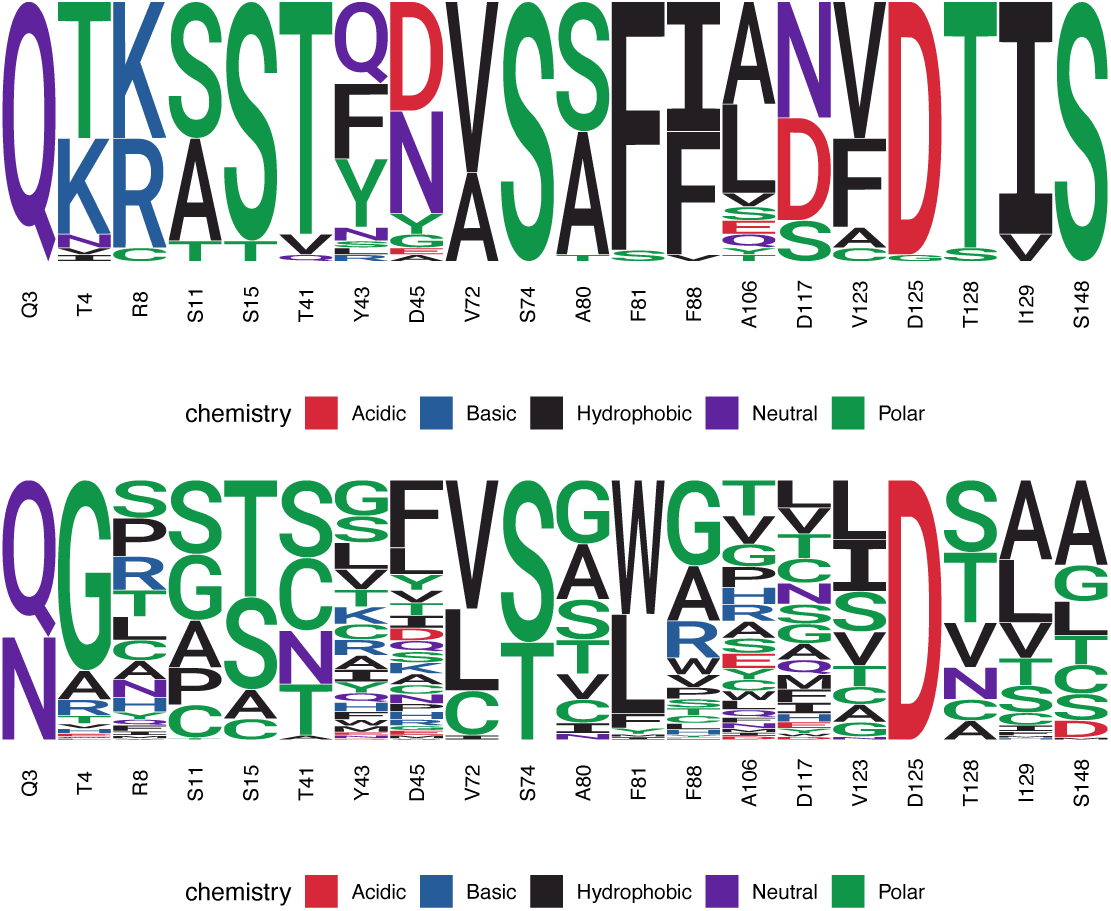
WebLogo plots of the 21 sites analysed in this study showing (A) phylogenetic conservation across microviridae (4 α3-like, 16 G4-like, and 18 ΦX174-like) and (B) mutational tolerance (this study). The height of the letters show the allele frequency among the 38 taxa (A) and the frequency of recovered plaques (B).

**Table 2.** Comparing the performance of three different strategies for predicting viability. Fit of different logistic regression models based on AIC is shown. Models using FoldX (FX) were run without predictor error included; PyRosetta (Py) estimates lack error estimates. dWeight and dHydrophobicity are the differences in amino acid molecular weight and hydrophobicity from the wildtype to the mutated residue. The rate4site conservation metric provides the best model for predicting viability. This metric requires a sequence alignment. We used 38 gene G sequences from microviridae (Figure 4).

| Model | AIC:<br>With (w/o error) |
| --- | --- |
| Blosum62 + dWeight +<br>dHydrophobicity | 524 |
| Py fold + FX trimer | 488 |
| rate4site | 428 |

**Figure 5.**
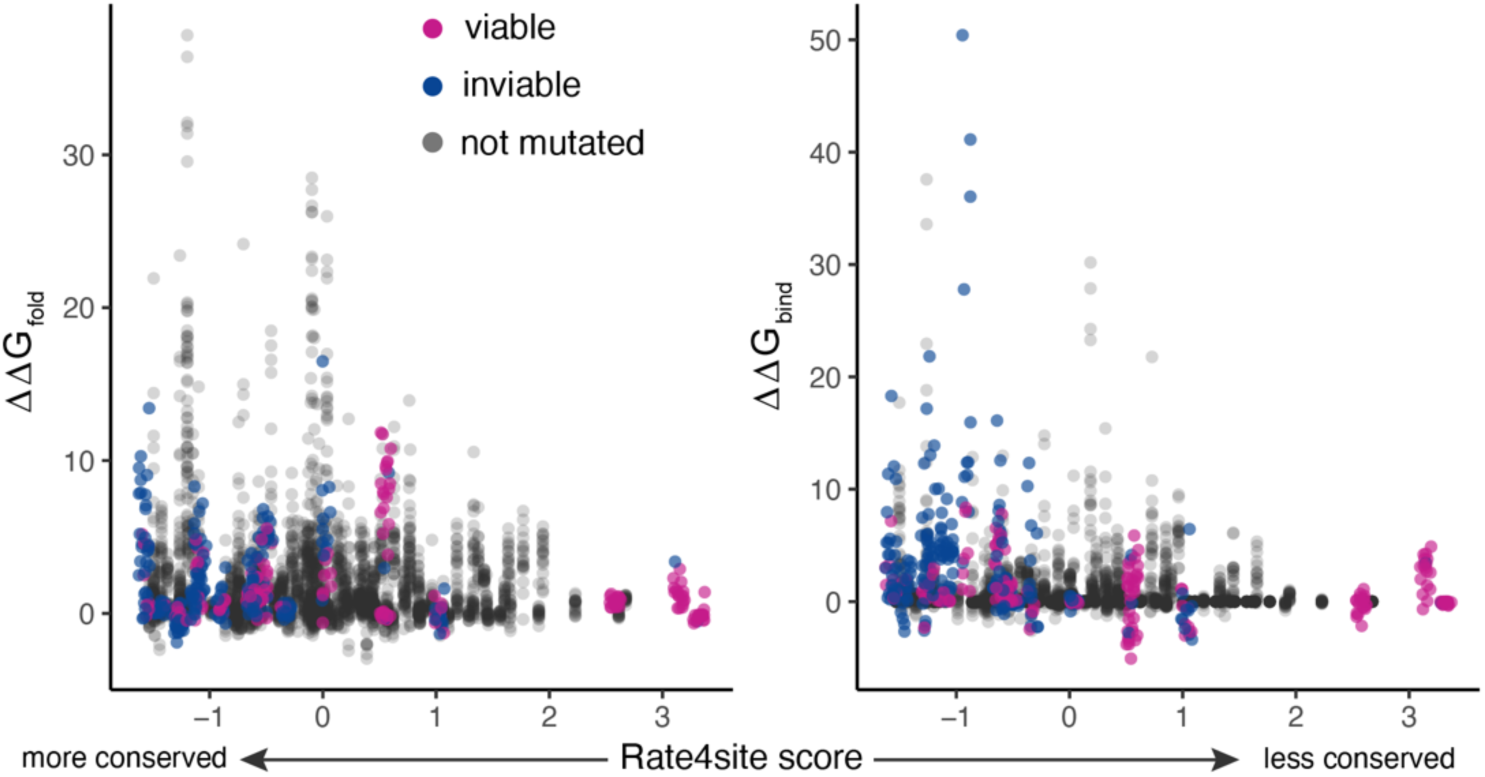
A comparison between phylogenetic conservation and predicted stability effects shows that variable residues (higher rate4site scores) always have low predicted effects on protein stability. The alternative is not true, highly conserved sites (low rate4site scores) often have some substitutions with high predicted effects on protein stability. Viable and inviable substitutions are shown in purple and blue, respectively.

### Synonymous codon biases

While we were primarily interested in modeling amino acid substitutions, we did construct all possible codons for each amino acid and noticed that some synonymous codons were recovered more or less often than expected based on their starting frequency in the initial library (**Supplementary figures 11-12, Supplementary data**). Thirteen codons were observed less frequently than expected based on the initial starting frequencies (p<0.01, see methods for description of multinomial model). Mutations observed less than expected were CGC-8-GTG, TCA-11-ACA, TCT-15-TGT, TCT-15-GCA, and ACG-41-GCG. Mutations observed more than expected were CAG-3-CAA, GAC-125-GAG, and ATT-129-CTC. Interestingly, 5/13 codons (TCT-15-TCT, GTT-72-GTT, TCT-74-TCT, TTT-81-TTT, TCT-148-TCT) that differed significantly from expected frequencies were wildtype codons, which were all observed less than expected. It has been observed that non-optimal codons are in some cases beneficial (Plotkin and Kudla 2010; Lyu and Liu 2020). We plan to follow up on this observation in future work on ΦX174. Of the 18 amino acids with unexpected observation frequencies (multinomial model, p<0.01), eight include a codon that is observed at unexpected frequencies. This led us to conclude that codon bias might have some influence on the plating efficiency of some amino acids, but is unlikely to impact whether or not an amino acid variant is viable.

### Plaque size measurements

We measured the plaque size of all 614 viable codon variants to 1) quantify another phenotype that could be related to viability and 2) because we were concerned about introducing biases in our plaque picking by accidentally missing very small plaques. Indeed, we found that some infrequently observed codon substitutions had isolates with significantly reduced plaque sizes (reported above in the “Defining viable versus inviable” section). This result caused us to increase our sampling efforts for some residues. Of the 614 codon variants measured, we found 156 with reduced plaque sizes (p<0.05, t-test, **Supplementary data**) with the average significantly smaller plaque size having a radius 68% smaller than wildtype. We found a weak but significant correlation between plaque size and plaque observation frequency (linear model, p=1.2E-12, r^2^=0.079). While most genotypes are observed about at their expected frequencies, genotypes that result in small plaques (less than about half the wildtype radius) are never picked at higher than expected frequencies (**Supplementary figure S13**). This is not true for large plaques, which occasionally are picked up to four times more often than expected.

## Discussion

It is challenging to predict the effects of mutations on protein function and consequently on the phenotypic characteristics of organisms making those proteins. However, progress towards improving these predictions has proceeded at a rapid pace (Wei and Li 2023). Our study contributes to this effort by 1) utilizing a study system of intermediate complexity with known protein binding partners, 2) performing extensive molecular modeling on the protein that we mutated and its binding partner, and 3) by manually building a large set of variants and measuring different phenotypes. We chose the G protein of ΦX174 because we expected this protein to be a good model for exploring how protein stability modeling can be mapped to viability because the G protein is a simple structural viral protein. It has no enzymatic activity and little host binding interaction. We performed some molecular modeling prior to building mutants and found wide variation in the predicted protein stability effects of amino acid substitutions (**Supplementary figure S8**). We chose to mutate residues that reflect this variation. Some mutations were predicted to have large effects, some small, some to affect binding, some to affect folding.

We found that ΦX174 tolerates about 50% of all possible single amino acid substitutions across the sites that we mutated. This is within the range (20-75%) found by others (Rennell et al. 1991; Markiewicz et al. 1994; Shafikhani et al. 1997; Guo et al. 2004). Some residues (e.g., Q3, D125) could not tolerate any substitutions, while others (D45, G119) could be mutated to any other amino acid. Across all 21 residues, we found that ΦX174 tolerates much greater amino acid variation than what is observed in microviridae sequences collected from nature (**Figure 4**). This suggests that many of the mutants we made may not be tolerated in natural conditions where there may be additional selective pressures impacting fitness. This finding supports the importance of the environment in genotype-phenotype mapping (Alberch 1991; Taverna and Goldstein 2002; Smith and Kruglyak 2008; Pigliucci 2010; Boucher et al. 2016; Moore et al. 2019). Other studies that performed mutational scanning of bacteriophages primarily targeted the receptor binding proteins (Andrews and Fields 2021; Huss et al. 2021). For example, Huss et al. (2021) focused on the receptor-binding portion of the tail fiber protein of T7 phage (Huss et al. 2021). They found a good correlation between ΔΔG predictions (Rosetta) and substitution tolerability, with 93% of the tolerated mutations predicted to be stabilizing. Thus the authors suggest that ΔΔG information should be used during protein engineering and that unexpected outcomes (e.g., stabilizing mutation isn’t tolerated) may point towards interesting biology. Our results differ from Huss et al. (2021) in that we found many destabilizing mutations to result in viable phages. However, the ΦX174 G protein modeling predicts most mutations to be destabilizing, so our ΔΔG distributions for ΦX174 are much different from the T7 tail fiber. It could be that our study included residues on the interior of the protein where Huss et al. (2021) focused on surface residues. It would be particularly interesting to investigate the evolutionary recovery of mutant constructs that we generated in this study. This approach often provides insight into how deleterious mutations cause fitness reductions. For example, Doore and Fane found that deleterious gene swaps between ΦX174 and G4 were ameliorated by beneficial mutations in scaffolding proteins (Doore and Fane 2015). They predicted that these scaffolding proteins improved protein folding and capsid assembly.

The three phenotypes (viability, observation frequency, plaque size) that we measured could only be partially predicted by amino acid properties, phylogenetic conservation, or molecular modeling. Molecular modeling did outperform basic amino acid properties (**Table 2**) but not sequence conservation information. However, these two best performing predictive methods are not very comparable because molecular modeling only requires data for the protein of interest while conservation information requires sequences from other taxa (preferably a good sampling of taxa). A main driver or error in the molecular modeling predictions arise from mutations that are inviable despite having low predicted folding effects. These types of mutations are common in our dataset and cause our logistic regression model relating viability to ΔΔG to asymptote at ∼0.7. Our sequence conservation-based metrics suffer from a lack of sequence diversity. There are many instances when little diversity is observed at a residue in the multiple sequence alignment, but the residue ends up tolerating mutations fairly well (residues T41, A80, F88, I129). It is these predictions that molecular modeling can do well with. Even though we were limited to a smaller number of taxa for measuring conservation, our results are similar to others. Høie et al. analyzed published data from 29 proteins (Høie et al. 2022). They correlated various sequence and molecular predictors of mutability to the results of deep mutational scanning and found high variation in the correlation between different proteins, with the strongest relationships being for the protein β-lactamase. Like in our work, Høie et al. (2022) found that many (∼25%) mutations predicted to have small effects on stability were inactive and that some of these errors could be remedied using sequence conservation metrics. A finding shared between Høie et al. (2022) and our work is that highly variable residues have no mutations that are predicted to change protein stability (**Figure 5**). The relationship between the conservation of a site and the predicted stability effects of mutations on that site suggests that amino acid variability is tolerated because the sites do not matter to protein stability. This means that the sequence diversity observed in ΦX174 is largely determined by protein stability and not other factors like escape from host defenses or adaptation to new host receptors. Our analysis of comparing the utility of molecular modeling versus phylogenetic conservation in predicting the outcome of mutations is confounded by the interdependency of these two things. The conservation of any given site and the average ΔΔG of folding and binding are correlated (**Figure 5**). The importance of the physical properties of proteins like solvent accessibility and secondary structure are evident in evolutionary conservation of these features (Ramsey et al. 2011). Protein physics is an important determinant of evolution (reviewed in (Bastolla et al. 2017)).

## Methods

### Molecular dynamics combined with FoldX

#### Structure file preparation

Crystal structure of the mature ΦX174 capsid was downloaded from the PDB using the 2BPA accession number (McKenna et al. 1992). The downloaded PDB file was modified to remove everything except pentamers of G and F capsid proteins i.e. five monomers each of G and F proteins (see **Figure 2**). The WHAT IF web server (https://swift.cmbi.umcn.nl) was then used to add missing atoms to the G-F protein complex structure.

#### System preparation and molecular dynamics simulations

A complete PDB file with G-F protein complex structure was used to prepare following five initial protein structures for carrying out atomistic molecular dynamics (MD) simulations: (i) a monomer of G protein; (ii) a dimer of G protein; (iii) a trimer of G protein; (iv) a pentamer of G protein; and (v) a complex of G and F protein pentamers. All five protein systems were subjected to MD simulations using the protocol reported in our previous studies (Miller et al. 2016; Patel et al. 2019). Briefly, the software package GROMACS 5.0.7 (Van Der Spoel et al. 2005) was used for all MD simulations with the Charmm22* forcefield (Chen et al. 2006). All five simulations were run for 100 ns and the simulation snapshots were saved every 1 ns resulting in 100 snapshots for each system.

#### FoldX analysis

Analysis of all the snapshots from five protein systems was carried out using FoldX software (Guerois et al. 2002; Schymkowitz et al. 2005). Similar to our prior studies (Miller et al. 2016; Patel et al. 2019), we first executed RepairPDB command using each snapshot as an input six times in succession to allow minimization of potential energy and to obtain its convergence.

Each energy minimized snapshot was then processed through BuildModel command to generate all possible 19 single mutations at each amino acid site of G protein in all five systems. Finally, folding stabilities (ΔΔ*G*_Fold_) were determined using Stability command on the monomer structures of G-protein and binding stabilities (ΔΔ*G*_Bind_) were estimated using AnalyseComplex command applied to the structures of the protein complexes (ii) to (v) mentioned above. For each mutation, we then calculated ΔΔ*G*_Fold_ and ΔΔ*G*_Bind_ values by averaging the FoldX estimates across all individual snapshot estimates.

To calculate ΔΔ*G*_Fold_ values for all possible 19 mutations at each amino acid site of the monomer of G protein, we carried out 332,500 FoldX calculations (175 G protein monomer residues × 19 possible mutations at each site × 100 MD snapshots). Whereas to calculate ΔΔ*G*_Bind_ values for protein complexes (ii) to (v) listed above, we performed: (1) 665,000 FoldX calculations for a dimer of G protein (175 G protein monomer residues × 19 possible mutations at each site × 100 MD snapshots × 2 copies of G monomers); (2) 997,500 FoldX calculations for a trimer of G protein (175 G protein monomer residues × 19 possible mutations at each site × 100 MD snapshots × 3 copies of G monomers); (3) 1,662,500 FoldX calculations for a pentamer of G protein (175 G protein monomer residues × 19 possible mutations at each site × 100 MD snapshots × 5 copies of G monomers); and (4) 332,500 FoldX calculations for a complex of G and F protein pentamers (175 G protein monomer residues × 19 possible mutations at each site × 100 MD snapshots). Averaging estimates across all individual snapshots ultimately resulted in 3,325 ΔΔ*G*_Fold_ and ΔΔ*G*_Bind_ values for each protein system (i) to (v) listed above. (see SI File)

#### PyRosetta

PyRosetta (Chaudhury et al. 2010) is a Python-based implementation of the Rosetta biomolecular modeling package (Rohl et al. 2004) that makes use of the Rosetta Monte Carlo sampling routines and scoring functions for protein folding (Ovchinnikov et al. 2018) and protein and small-molecule docking (Chaudhury and Gray 2008; Davis and Baker 2009). PyRosetta v4.0 (Chaudhury et al. 2010) was used to sample conformational changes to the wild type G protein of ΦX174 (2BPA) upon mutation and calculate its folding and binding free energy differences, ΔΔ*G*_Fold_ and ΔΔ*G*_Bind_, respectively.

#### Folding

To determine ΦX174 folding free energy changes, we first relaxed the wild type protein structure by repacking all of its side chains by sampling from the 2010 Dunbrack rotamer Library (Shapovalov and Dunbrack 2011). This sampling employs a Monte Carlo search routine that iteratively swaps rotamers of a randomly-selected residue from the protein structure and accepts/rejects the swap according to the Metropolis criterion (Metropolis et al. 1953). We then minimized the wild type structure via a linear minimization scheme using the scoring function from (Alford et al. 2017). After relaxing the protein structure, each missense mutation was sequentially introduced and all residues within a 10 Å distance of the mutated residue’s center were repacked, followed by a Monte Carlo sampling coupled with energy minimization of the backbone and side-chain conformation to optimize the protein geometry (Firnberg et al. 2014). To ensure folding free energy convergence, each simulation consisted of 300,000 Monte Carlo iterations. We obtained ΔΔ*G*_Fold_ values for all possible 19 mutations at the 21 empirically mutated sites of the monomer of G protein.

#### Binding

We employed the PyRosetta Docking Protocol (Chaudhury et al. 2011) to predict the binding free energies of the G protein dimers and trimers. The protocol consists of a perturbation stage and a docking stage. To quickly identify favorable docking poses, we started the perturbation stage by first converting the complex into a centroid form. The perturbation stage consists of global perturbation, which introduces a random initial orientation of the two binding partners, translational perturbations, which translate one docking partner away from the other in a random direction and at a random distance, and rotational perturbations, which rotate one binding partner around an axis at a random angle. Because proteins are not spherical, sometimes the random orientations result in severe clashes between the binding partners, and at other times, the binding partners are placed so far apart that they no longer touch. Hence, as part of our binding protocol, we then translated the downstream binding protein along the line of protein centers until the two binding partners are in contact. Next, we converted the complex back to all-atom form and recovered the original side-chain conformations to minimize on the docking jump. The docking stage employs a multi-scale Monte Carlo based docking algorithm that incorporates both a low-resolution, centroid-mode docking and a high resolution, all-atom mode docking (Firnberg et al. 2014). The low-resolution docking can quickly locate binding interfaces, and the high-resolution docking can optimize rigid-body orientation and side-chain conformation. Once the low-resolution docking is finished, the lowest energy structure is selected for high-resolution docking refinement. For exhaustive searches with PyRosetta docking, for every mutant, we generated 500 decoys or candidate structures. We assessed docking performance by sorting these respective decoy sets by the total docking score and the complex with the best score was selected as the final structural model for computing ΔΔ*G*_Bind._

### Viral mutagenesis and propagation

#### Generating variants

We used a 2-step PCR reaction to perform site-directed mutagenesis on the laboratory ΦX174 strain (Genbank Accession AF176034). Briefly, the viral genome was amplified in two approximately equal-length products with degenerate primers containing NNN at the codon to be mutagenized. The ends of these ½ genome fragments contain overlapping complementary sticky end that used to anneal the ½ genome fragments together in a second two-step thermocycling reaction (Van Leuven et al. 2021). The circularized amplicons were transformed into electrocompetent *E. coli* C cells and plated on ΦLB plates (10g/L tryptone, 5g/L Bacto yeast extract, 10g/L NaCl, 2 mM CaCl_2_) in 0.3% ΦLB top agar. Plates were incubated 4-5 hours at 37°C. The plaque of each viable phage was scraped off and stored individually in 200 uL of ΦLB without CaCl_2_ in a deep 96-well plate.

Initially, 94 of the phage harvested from each mutagenesis were PCR amplified with phiX0F and 2953R primers and the DNA sequenced across the codon of interest by Eurofins Genomics (Louisville, KY) or locally on an ABI 3100 using BigDye terminator chemistry (ThermoFisher). Further replication and final sample size issues are discussed under *Viability Analysis* below. The numbers of each codon recovered were compiled using R. Briefly, the sangerseqR package was used to read and process all of the Sanger reads by trimming 100 bases off the 5’ end and keeping the following 400 bp for analysis. Sequences with more than three mismatches to the G gene across this 300 bp window were discarded. The major and secondary peaks were identified and we required a maximum secondary peak ratio of 0.25 at the mutated codon.

Around ∼10% of sequences exceeded this ratio. These plaques contained at least two distinct virus genotypes and were discarded. We considered these plaques to result from co-transfected host cells which could alter our viability analysis because an inviable virus could be complemented by a co-transfected viable virus. Most of the Sanger reads were checked manually in Lasergene.

#### Illumina sequencing

Viral DNA was amplified using forward primer (5’-ACACTGACGACATGGTTCTACAGTCGCGGTAGGTTTTCTGCTTA-3’) and reverse primer (5’-TACGGTAGCAGAGACTTGGTCTCCACCAGCAAGAGCAGAAGCA-3’) annealing to bases 2364 and 2975, respectively. PCR products were purified using magnetic beads and quantified by gel electrophoresis into 5 quantity bins. These discrete classifications were used to determine the amount of 1st round PCR products used for the 2^nd^ round barcoding PCR step (U’Ren and Arnold 2022). This approximate normalization procedure equalizes the concentration of 2^nd^ round PCR products. Reads were mapped to the ΦX174 G gene using bwa-mem with modified clipping and mismatch penalties (-L 20 -B 1) to account for mutagenesis at the 5’ end of the genes. A minimum of 10 reads with mapping quality greater than 50 were required for calling variant codons. Codon counts were parsed from BAM files using the R package Rsamtools. In our first attempt, sites 4 and 125 displayed severe non-uniform coverage of each of the 64 codons in the starting pool. We re-ordered these mutagenesis primers from another supplier (IDT) and sequenced another 86 and 82 plaques, respectively. At both sites we observed additional viable amino acid variants and thus decided to use the data using the IDT primers, although data for both are included in this paper. An arbitrary cutoff of 10 reads was required for the identification of codon variants.

#### Measuring plaque size

After Sanger sequencing, one plaque for every codon was chosen from the 200 uL ΦLB stocks and replated on ΦLB plates with log-phase *E. coli* C hosts. We ensured that 30-150 plaques were on the plate to be measured by altering the dilution if needed. Plate images were collected using a Canon DSLR camera with a stand and the images were analyzed in OpenCFU. The camera to plate distance was kept constant to not introduce differences between plate images. OpenCFU plaque size counts were compiled and analyzed in R. Plaque size distributions often have a long tail of small-sized plaques. The 90% quantile of plaque radiuses was used to avoid the influence of this tail. We used prop.test in R to compare the ratio of small or normal sized plaques between different groups of variants (e.g., amino acids observed at lower then expected frequency). Multiple t-tests with Bonferroni p-value correction were performed in R to determine if plaque sizes for a variant were smaller than wildtype.

#### Bioinformatics

Amino acid conservation was calculated using rate4site using alignments compiled from all available G protein sequences. These sequences were attained by using BlastP to search nr for G protein sequences (blastp -evalue 1E-10 -query NP_040712.1 -db nr). The resulting list of protein sequences was curated by removing sequences that were shorter than 131 amino acids in length (75% of ΦX174) and collapsing sequences that were greater than 99% identical using cdhit v4.8.4 (-c 0.99). The resulting 78 amino acid sequences were aligned using Muscle v3.8 (default parameters). Per-site conservation values were calculated using rate4site (v2.01) in Jalview using the AMAS method described in (Livingstone and Barton 1993). SIFT values were accessed through the web server and run with default parameters (Kumar et al. 2009).

Protein surface exposure was determined using the Accessible Surface Area and Accessibility calculator from (Anon) and the dssp calculation from (Kabsch and Sander 1983) using the G pentamer pdb file. Surface and buried designations as shown in **Supplementary figure S8** were determined using cutoffs, with surface residues having RSA values greater than 0.3 and buried residues values less than 0.05. The location of b-sheets in the protein secondary structure was determined from UniProt. Residues involved in subunit interactions were identified from (McKenna et al. 1994).

#### Viability Analysis

Amino acid variants observed once or more through sequencing plaques are known to be viable; unobserved variants may be inviable or they may be viable but unsampled. We used the multinomial model within a Bayesian approach to determine the probability that an unobserved mutation is inviable. For each site we estimated the frequency of each amino acid prior to transformation by sequencing the ligation mix (see *Illumina sequencing methods* above). For two sites (4 and 125), sequencing revealed codon frequencies were extremely uneven; for these two sites, we ordered new mixes and here only include data from the replacement mixes. Most ligation mixes were sequenced a single time; for sites 3, 4, 11, and 125, however, we sequenced three samples from each mix and then summed read counts across triplicate samples. At each site, codon-read counts were then pooled to obtain amino acid frequencies (*p*). We next assumed that amino acid level count data come from a multinomial sample of the ligation mix where inviable mutations have probability zero and all other mutations are sampled with probability equal to their relative frequency among the set of viable amino acids. Below, under *Plating Efficiency*, we address the validity of this assumption. Let the set of viable (1) and inviable (0) amino acids at a site be summarized as a 20-element binary vector, *B_i_*, where *i* indexes a particular 0/1 combination (*i* = 1, 2, 3…2^20^). Our prior was that all 2^20^-1 binary vectors (excluding 00000000000000000000) are equally likely. For a given site, we then calculated the multinomial probability of the observed vector of count, C, given *B_i_* (and *p*) and did so for all *i*. We assumed *p* was known without error from the ligation-mix sequencing. The posterior probability of any particular binary vector, *B_i_*, comes from application of Bayes’ formula:

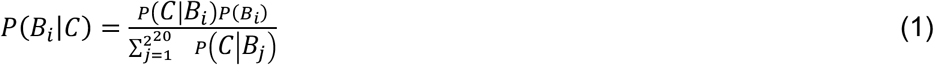

where *P*(*C*|*B*_*i*_) follows the multinomial distribution with category probabilities *p*. Note that for many *B_i_*, the P(*B_i_*|C)=0 because *B_i_* contains a zero at one or more amino acids that appear in *C*; for computational efficiency, we need only do this calculation over the subset of *B* with probability > 0 (i.e. the subset of vectors consistent with the data). One important thing that comes from eq 1 is the ability to calculate the posterior probability of the particular *B_i_* with 1s that exactly match the observed amino acids and 0s at the unobserved ones–i.e. the probability that the observed viability vector is the true viability vector, or P_obs=true_. When P_obs=true_ is close to one, our model indicates further replication is not necessary. The second important calculation is the probability that a particular unobserved amino acid, call it *X*, is viable: P(*X*=1|C). This is obtained by summing the posterior probability of all vectors within *B* where *X*=1 using the indicator function *I*(*X∊B_i_*)that takes value 1 when *X=1* within *B_j_*, and 0 otherwise:

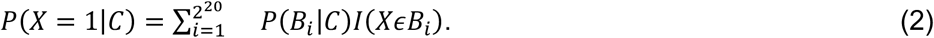

In practice, we initially sequenced 94 plaques at each site (not all of which yielded usable data). We then used eq 1 to calculate P_obs=true_. When P_obs=true_ for a site was < 0.95, we sequenced more plaques (usually 46 more), returned to eq 1. If necessary, we iterated a third or fourth time. After sequencing an average of 140 plaques per residue (range 89-204) we stopped to reevaluate. At the 10 sites (3, 8, 15, 41, 43, 45, 74, 106, 117, and 125) where P_obs=true_ exceeded 0.99 we stopped for good and assumed unobserved amino acids were inviable. At the other sites P_obs=true_ was < 0.99. This occurred when one or more amino acids were very rare in a ligation mix (i.e. when an amino acid is very rare in a mix, the failure to observe it says little about viability). At these sites we used eq 2 to pinpoint 31 particular site/amino-acid combinations might be viable based on having posteriors > 0.1%. For these, we designed (degenerate) primers to synthesize the variant with only the target amino acid. We attempted to make each variant three times. When triplicate attempts yielded no plaques, we designated the variant as inviable. Plaques were sequenced to confirm the identity of the variant. After this work was completed, we discovered one oversight: W at site 128 has an 11% posterior probability of being viable but we failed to synthesize it. We have excluded this variant from the analysis.

#### Plating Efficiency

The viability analysis assumes that viable variants are observed in proportion to their frequency in the ligation mix. This assumption is supported by the fact that, among known viable variants, the correlation between ligation-mix-based expected counts and observed counts is high: r=0.91. See Supplemental Materials for scatterplots of expected vs observed for all sites pooled and for sites individually. However, when we simulate 1000 multinomial samples using the ligation mix frequencies and our real sample sizes, we observe an average r=0.97 and no simulated r < 0.95. Thus, some variants are observed more often or less often than their ligation mix frequency would suggest. The simple explanation for this is that some variants exert strong effects on plating efficiency. Plating efficiency is the term we use to describe how, when a cell is infected (in this case, via transformation), different variants can differ in the probability of forming a plaque. We sought to identify which variants showed significant plating efficiency effects using the following stepwise likelihood ratio test framework applied to each site. The null hypothesis was that the observed counts arose as a multinomial sample from the ligation-mix amino acid frequencies (normalized to the viable amino acids). The first alternative was that all amino acids are sampled according to the ligation mix except one; this one amino acid instead has a multinomial sampling probability estimated from its observed count frequency. We calculated the multinomial log-likelihood (L) under the null and the alternative and defined their difference to be Λ_real_ = L_alt_ - L_null_. In practice, we considered each amino acid as the potential first one removed to constitute the alternative; we used whichever one maximized Λ. To test if the log-likelihood improvement was significant, we then simulated 1000 datasets using the frequencies assumed under the null and, for each, ran through the process of calculating Λ. Denoting these 1000 values as Λ_sim_, we then estimated the p-value as the proportion of Λ_sim_ ≥ Λ_real_. We then continued in a stepwise fashion: the step-one alternative was defined as the null and we considered a second-step alternative where a second amino acid takes a multinomial frequency from the observed counts instead of the ligation mix. The algorithm was continued until all but one viable amino acid had their sampling frequency reassigned. The p-value in this process increases monotonically; if the step-one alternative hypothesis is not significant, then the site is consistent with the null.

### Multistage binomial model

The multistage binomial (MSB) model describes a system in which a binary outcome results from multistage processes with multiplicative risks. Its basic construct is as follows:

In a system of *n* units, the binary outcome *Y*_*i*_ of a process with *q* stages and covariate vector ***W***_*ij*_ (*j* = 1,2,…,*q*) is given as:

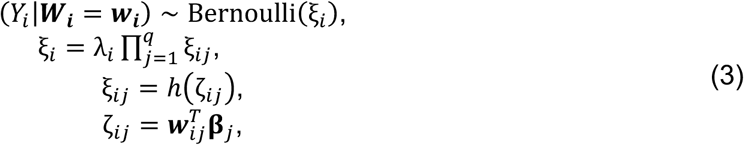

where ξ_*i*_ denotes success probability, λ_*i*_ is the upper limit of ξ_*i*_ for each Bernoulli trial given ***W***_*i*_, ℎ(ζ_*ij*_) = 1/*1 + *e*^-ζ_ij_^) is the inverse link function, and **β**_*j*_ is a *p*_*j*_-vector of regression coefficients at stage *j*. In this model, each stage in the process is an independent Bernoulli experiment with the final outcome corresponding to all experiments being successful. The success probability ξ_*i*_ is bounded by [0, λ_*i*_], which represents theoretical limits that may arise from unknown stages or known stages with no measured explanatory variable. Model (3) can be generalized to cases with error-prone covariates in which the covariate matrix ***X***_***i***_ is such that ***X***_***ij***_ = ***W***_***ij***_ + ***e***_***ij***_ where ***e***_***ij***_ is a measurement error vector. The parameters are then estimated using penalized maximum likelihood. The details of the MSB model and the estimation process are described in (Tovissodé et al. 2025).

## Author Contributions

JTVL, HAW, FMY, BMR, CRM conceived and planned the experiments. JTVL, LS, KB, EA carried out the experiments. JSP, JY, CB performed the molecular modeling. YS, CFT, CRM developled and implemented the MBP model. JTVL, HAW, BMR, CRM contributed to the interpretation of the results and statistical analysis. JTVL, CRM, BMR contributed to writing the manuscript. All authors provided critical feedback and helped shape the research, analysis and manuscript.

## Acknowledgments

This study was supported by the National Institutes of Health (NIH) grants P20 GM104420 and P20 GM103397 and by the National Science Foundation (NSF) grant EPSCoR Track-II, Award No. OIA-1736253. We thank the Institute for Bioinformatic and Evolutionary Studies (IBEST) Genomics Resources Core for assistance with DNA sequencing library preparation.

## Data Availability

Code and data are available on GitHub at <u>yanghaobojordan/PhiX</u> and <u>jtvanleuven/phiX_biophysical_modeling</u>. Raw Illumina sequencing data has been deposited in GenBank under accession numbers SRAXXXX-SRAXXXXX.

## Supplementary data

Spreadsheet containing useful raw data and results too large to fit into supplementary tables. Sheets “obs_vs_exp_AA” and “obs_vs_exp_codon” are plaque observation counts, frequencies (normalized to sum), expected frequencies (based on Illumina sequencing of initial pools), expected versus observed statistics. The “starting_read_counts_by_codon” sheet contains the Illumina read counts for each codon in the initial phage pool (see Methods for details). These numbers include the number of reads for each library and the number of reads mapped to the G gene sequence. Codon counts were parsed from *.bam alignment files. The “molecular_modeling” sheet contains all of the FoldX (FX) and PyRosetta (Py) ΔΔG predictions for folding (of monomers) and binding (see Figure 1 for description of different binding simulations). The “plaque_size” sheet contain<u>s</u> the summarized results from OpenCFU. For every genotype, one plaque was chosen (“plaque” number), amplified, and plated. The “single” column indicates that this codon was generated specifically by site-directed mutagenesis instead of the recovered from the library. All of the plate pictures, analysis code, and non-summarized plaque size data are available at <u>jtvanleuven/phiX_biophysical_modeling</u> in the “rawdata/plate_images” folder. The “rate4site” and “SIFT” sheets contain the output of these programs.

## References

Adzhubei IA, Schmidt S, Peshkin L, Ramensky VE, Gerasimova A, Bork P, Kondrashov AS, Sunyaev SR. 2010. A method and server for predicting damaging missense mutations. Nat. Methods 7:248–249.

Alberch P. 1991. From Genes to Phenotype - Dynamic-Systems and Evolvability. Genetica 84:5–11.

Alford RF, Leaver-Fay A, Jeliazkov JR, O’Meara MJ, DiMaio FP, Park H, Shapovalov MV, Renfrew PD, Mulligan VK, Kappel K, et al. 2017. The Rosetta All-Atom Energy Function for Macromolecular Modeling and Design. J. Chem. Theory Comput. 13:3031–3048.

Andrews B, Fields S. 2021. Balance between promiscuity and specificity in phage λ host range. ISME J. 15:2195–2205.

Anon. Calculate accessible surface area and accessibility of protein. Available from: http://cib.cf.ocha.ac.jp/bitool/ASA/

Barnes JE, Miller CR, Ytreberg FM. 2022. Searching for a mechanistic description of pairwise epistasis in protein systems. Proteins Struct. Funct. Bioinforma. [Internet] n/a. Available from: https://onlinelibrary.wiley.com/doi/abs/10.1002/prot.26328

Bastolla U, Dehouck Y, Echave J. 2017. What evolution tells us about protein physics, and protein physics tells us about evolution. Curr. Opin. Struct. Biol. 42:59–66.

Beaulieu JM, O’Meara BC, Zaretzki R, Landerer C, Chai J, Gilchrist MA. 2019. Population Genetics Based Phylogenetics Under Stabilizing Selection for an Optimal Amino Acid Sequence: A Nested Modeling Approach. Mol. Biol. Evol. 36:834–851.

Bloom JD. 2014. An Experimentally Determined Evolutionary Model Dramatically Improves Phylogenetic Fit. Mol. Biol. Evol. 31:1956–1978.

Bloom JD, Silberg JJ, Wilke CO, Drummond DA, Adami C, Arnold FH. 2005. Thermodynamic prediction of protein neutrality. Proc. Natl. Acad. Sci. 102:606–611.

Boucher JI, Bolon DNA, Tawfik DS. 2016. Quantifying and understanding the fitness effects of protein mutations: Laboratory versus nature. Protein Sci. 25:1219–1226.

Bull JJ, Heineman RH, Wilke CO. 2011. The Phenotype-Fitness Map in Experimental Evolution of Phages. PLOS ONE 6:e27796.

Chaudhury S, Berrondo M, Weitzner BD, Muthu P, Bergman H, Gray JJ. 2011. Benchmarking and Analysis of Protein Docking Performance in Rosetta v3.2. PLOS ONE 6:e22477.

Chaudhury S, Gray JJ. 2008. Conformer selection and induced fit in flexible backbone protein-protein docking using computational and NMR ensembles. J. Mol. Biol. 381:1068–1087.

Chaudhury S, Lyskov S, Gray JJ. 2010. PyRosetta: a script-based interface for implementing molecular modeling algorithms using Rosetta. Bioinformatics 26:689–691.

Chen J, Im W, Brooks CL. 2006. Balancing Solvation and Intramolecular Interactions: Toward a Consistent Generalized Born Force Field. J. Am. Chem. Soc. 128:3728–3736.

Crill WD, Wichman HA, Bull JJ. 2000. Evolutionary Reversals During Viral Adaptation to Alternating Hosts. Genetics 154:27–37.

Davis IW, Baker D. 2009. RosettaLigand docking with full ligand and receptor flexibility. J. Mol. Biol. 385:381–392.

DePristo MA, Weinreich DM, Hartl DL. 2005. Missense meanderings in sequence space: a biophysical view of protein evolution. Nat. Rev. Genet. 6:678–687.

Doore SM, Fane BA. 2015. The Kinetic and Thermodynamic Aftermath of Horizontal Gene Transfer Governs Evolutionary Recovery. Mol. Biol. Evol. 32:2571–2584.

Duan J, Lupyan D, Wang L. 2020. Improving the Accuracy of Protein Thermostability Predictions for Single Point Mutations. Biophys. J. 119:115–127.

Fares MA, Ruiz-González MX, Moya A, Elena SF, Barrio E. 2002. GroEL buffers against deleterious mutations. Nature 417:398–398.

Faure AJ, Martí-Aranda A, Hidalgo-Carcedo C, Beltran A, Schmiedel JM, Lehner B. 2024. The genetic architecture of protein stability. Nature 634:995–1003.

Firnberg E, Labonte JW, Gray JJ, Ostermeier M. 2014. A Comprehensive, High-Resolution Map of a Gene’s Fitness Landscape. Mol. Biol. Evol. 31:1581–1592.

Gonzalez CE, Ostermeier M. 2019. Pervasive Pairwise Intragenic Epistasis among Sequential Mutations in TEM-1 β-Lactamase. J. Mol. Biol. [Internet]. Available from: http://www.sciencedirect.com/science/article/pii/S0022283619301512

Grantham R. 1974. Amino Acid Difference Formula to Help Explain Protein Evolution. Science 185:862–864.

Guerois R, Nielsen JE, Serrano L. 2002. Predicting changes in the stability of proteins and protein complexes: a study of more than 1000 mutations. J. Mol. Biol. 320:369–387.

Guo HH, Choe J, Loeb LA. 2004. Protein tolerance to random amino acid change. Proc. Natl. Acad. Sci. 101:9205–9210.

Hill AM, Ingle TA, Wilke CO. 2024. A computational model for bacteriophage ϕX174 gene expression. PLOS ONE 19:e0313039.

Høie MH, Cagiada M, Frederiksen AHB, Stein A, Lindorff-Larsen K. 2022. Predicting and interpreting large-scale mutagenesis data using analyses of protein stability and conservation. Cell Rep. [Internet] 38. Available from: https://www.cell.com/cell-reports/abstract/S2211-1247(21)01711-3

Huss P, Meger A, Leander M, Nishikawa K, Raman S. 2021. Mapping the functional landscape of the receptor binding domain of T7 bacteriophage by deep mutational scanning. eLife 10:e63775.

Jack BR, Wilke CO. 2019. Pinetree: a step-wise gene expression simulator with codon-specific translation rates. Bioinformatics 35:4176–4178.

Kabsch W, Sander C. 1983. Dictionary of protein secondary structure: Pattern recognition of hydrogen-bonded and geometrical features. Biopolymers 22:2577–2637.

Kawashima S, Kanehisa M. 2000. AAindex: Amino Acid index database. Nucleic Acids Res. 28:374.

Koehler Leman J, Weitzner BD, Lewis SM, Adolf-Bryfogle J, Alam N, Alford RF, Aprahamian M, Baker D, Barlow KA, Barth P, et al. 2020. Macromolecular modeling and design in Rosetta: recent methods and frameworks. Nat. Methods 17:665–680.

Kumar P, Henikoff S, Ng PC. 2009. Predicting the effects of coding non-synonymous variants on protein function using the SIFT algorithm. Nat. Protoc. 4:1073–1081.

Lee KH, Miller CR, Nagel AC, Wichman HA, Joyce P, Ytreberg FM. 2011. First-Step Mutations for Adaptation at Elevated Temperature Increase Capsid Stability in a Virus. PLOS ONE 6:e25640.

Levy SF, Blundell JR, Venkataram S, Petrov DA, Fisher DS, Sherlock G. 2015. Quantitative evolutionary dynamics using high-resolution lineage tracking. Nature 519:181–186.

Li B, Yang YT, Capra JA, Gerstein MB. 2020. Predicting changes in protein thermodynamic stability upon point mutation with deep 3D convolutional neural networks. PLOS Comput. Biol. 16:e1008291.

Livingstone CD, Barton GJ. 1993. Protein sequence alignments: a strategy for the hierarchical analysis of residue conservation. Comput. Appl. Biosci. CABIOS 9:745–756.

Lyu X, Liu Y. 2020. Nonoptimal Codon Usage Is Critical for Protein Structure and Function of the Master General Amino Acid Control Regulator CPC-1. mBio 11:e02605–20.

Markiewicz P, Kleina LG, Cruz C, Ehret S, Miller JH. 1994. Genetic Studies of the lac Repressor. XIV. Analysis of 4000 Altered Escherichia coli lac Repressors Reveals Essential and Non-essential Residues, as well as “Spacers” which do not Require a Specific Sequence. J. Mol. Biol. 240:421–433.

McKenna R, Ilag LL, Rossmann MG. 1994. Analysis of the Single-stranded DNA Bacteriophage φX174, Refined at a Resolution of 3·0 Å. J. Mol. Biol. 237:517–543.

McKenna R, Xia D, Willingmann P, Ilag LL, Krishnaswamy S, Rossmann MG, Olson NH, Baker TS, Incardona NL. 1992. Atomic structure of single-stranded DNA bacteriophage ΦX174 and its functional implications. Nature 355:137–143.

Metropolis N, Rosenbluth AW, Rosenbluth MN, Teller AH, Teller E. 1953. Equation of State Calculations by Fast Computing Machines. J. Chem. Phys. 21:1087–1092.

Miller CR, Johnson EL, Burke AZ, Martin KP, Miura TA, Wichman HA, Brown CJ, Ytreberg FM. 2016. Initiating a watch list for Ebola virus antibody escape mutations. PeerJ 4:e1674.

Moore R, Casale FP, Jan Bonder M, Horta D, Franke L, Barroso I, Stegle O. 2019. A linear mixed-model approach to study multivariate gene–environment interactions. Nat. Genet. 51:180–186.

Ovchinnikov S, Park H, Kim DE, DiMaio F, Baker D. 2018. Protein structure prediction using Rosetta in CASP12. Proteins 86 Suppl 1:113–121.

Pancotti C, Benevenuta S, Birolo G, Alberini V, Repetto V, Sanavia T, Capriotti E, Fariselli P. 2022. Predicting protein stability changes upon single-point mutation: a thorough comparison of the available tools on a new dataset. Brief. Bioinform. 23:bbab555.

Patel JS, Quates CJ, Johnson EL, Ytreberg FM. 2019. Expanding the watch list for potential Ebola virus antibody escape mutations. PloS One 14:e0211093.

Pepin KM, Wichman HA. 2007. Variable epistatic effects between mutations at host recognition sites in phiX174 bacteriophage. Evol. Int. J. Org. Evol. 61:1710–1724.

Pigliucci M. 2010. Genotype–phenotype mapping and the end of the ‘genes as blueprint’ metaphor. Philos. Trans. R. Soc. B Biol. Sci. 365:557–566.

Plotkin JB, Kudla G. 2010. Synonymous but not the same: the causes and consequences of codon bias. Nat. Rev. Genet. 12:32–42.

Ramsey DC, Scherrer MP, Zhou T, Wilke CO. 2011. The Relationship Between Relative Solvent Accessibility and Evolutionary Rate in Protein Evolution. Genetics 188:479–488.

Rennell D, Bouvier SE, Hardy LW, Poteete AR. 1991. Systematic mutation of bacteriophage T4 lysozyme. J. Mol. Biol. 222:67–88.

Rohl CA, Strauss CEM, Misura KMS, Baker D. 2004. Protein Structure Prediction Using Rosetta. In: Methods in Enzymology. Vol. 383. Numerical Computer Methods, Part D. Academic Press. p. 66–93. Available from: http://www.sciencedirect.com/science/article/pii/S0076687904830040

Sapozhnikov Y, Patel JS, Ytreberg FM, Miller CR. 2023. Statistical modeling to quantify the uncertainty of FoldX-predicted protein folding and binding stability. BMC Bioinformatics 24:426.

Sarkisyan KS, Bolotin DA, Meer MV, Usmanova DR, Mishin AS, Sharonov GV, Ivankov DN, Bozhanova NG, Baranov MS, Soylemez O, et al. 2016. Local fitness landscape of the green fluorescent protein. Nature 533:397–401.

Schymkowitz J, Borg J, Stricher F, Nys R, Rousseau F, Serrano L. 2005. The FoldX web server: an online force field. Nucleic Acids Res. 33:W382–388.

Shafikhani S, Siegel RA, Ferrari E, Schellenberger V. 1997. Generation of Large Libraries of Random Mutants in Bacillus subtilis by PCR-Based Plasmid Multimerization. BioTechniques 23:304–310.

Shapovalov MV, Dunbrack RL. 2011. A Smoothed Backbone-Dependent Rotamer Library for Proteins Derived from Adaptive Kernel Density Estimates and Regressions. Structure 19:844–858.

Smith EN, Kruglyak L. 2008. Gene–Environment Interaction in Yeast Gene Expression. PLOS Biol. 6:e83.

Sun Y, Roznowski AP, Tokuda JM, Klose T, Mauney A, Pollack L, Fane BA, Rossmann MG. 2017. Structural changes of tailless bacteriophage ΦX174 during penetration of bacterial cell walls. Proc. Natl. Acad. Sci.:201716614.

Taverna DM, Goldstein RA. 2002. Why are proteins so robust to site mutations? J. Mol. Biol. 315:479–484.

Tokuriki N, Stricher F, Schymkowitz J, Serrano L, Tawfik DS. 2007. The Stability Effects of Protein Mutations Appear to be Universally Distributed. J. Mol. Biol. 369:1318–1332.

Tokuriki N, Tawfik DS. 2009. Chaperonin overexpression promotes genetic variation and enzyme evolution. Nature 459:668–673.

Tovissodé CF, Sapozhnikov Y, Miller C. 2025. A Multistage Binomial Model with Measurement Errors: Application to Protein Viability Prediction. Available from: https://www.preprints.org/manuscript/202509.2376

U’Ren JM, Arnold AE. 2022. Illumina MiSeq Dual-barcoded Two-step PCR Amplicon Sequencing Protocol. Available from: https://www.protocols.io/view/illumina-miseq-dual-barcoded-two-step-pcr-amplicon-b748rqzw

Vale PF, Choisy M, Froissart R, Sanjuán R, Gandon S. 2012. The Distribution of Mutational Fitness Effects of Phage φX174 on Different Hosts. Evolution 66:3495–3507.

Van Der Spoel D, Lindahl E, Hess B, Groenhof G, Mark AE, Berendsen HJC. 2005. GROMACS: Fast, flexible, and free. J. Comput. Chem. 26:1701–1718.

Van Leuven JT, Ederer MM, Burleigh K, Scott L, Hughes RA, Codrea V, Ellington AD, Wichman HA, Miller CR. 2021. ΦX174 Attenuation by Whole-Genome Codon Deoptimization. Genome Biol. Evol. [Internet] 13. Available from: 10.1093/gbe/evaa214

Venkataram S, Dunn B, Li Y, Agarwala A, Chang J, Ebel ER, Geiler-Samerotte K, Hérissant L, Blundell JR, Levy SF, et al. 2016. Development of a Comprehensive Genotype-to-Fitness Map of Adaptation-Driving Mutations in Yeast. Cell 166:1585–1596.e22.

Wei H, Li X. 2023. Deep mutational scanning: A versatile tool in systematically mapping genotypes to phenotypes. Front. Genet. [Internet] 14. Available from: https://www.frontiersin.org/journals/genetics/articles/10.3389/fgene.2023.1087267/full

Weinreich DM, Watson RA, Chao L. 2005. Perspective: Sign Epistasis and Genetic Constraint on Evolutionary Trajectories. Evolution 59:1165–1174.

Wylie CS, Shakhnovich EI. 2011. A biophysical protein folding model accounts for most mutational fitness effects in viruses. Proc. Natl. Acad. Sci. 108:9916–9921.

Yang J, Naik N, Patel JS, Wylie CS, Gu W, Huang J, Ytreberg FM, Naik MT, Weinreich DM, Rubenstein BM. 2020. Predicting the viability of beta-lactamase: How folding and binding free energies correlate with beta-lactamase fitness. PLOS ONE 15:e0233509.

