## supplementary_figures for "ΦX174 bacteriophage viability predicted by protein biophysical modeling"

### Supplementary figures and tables

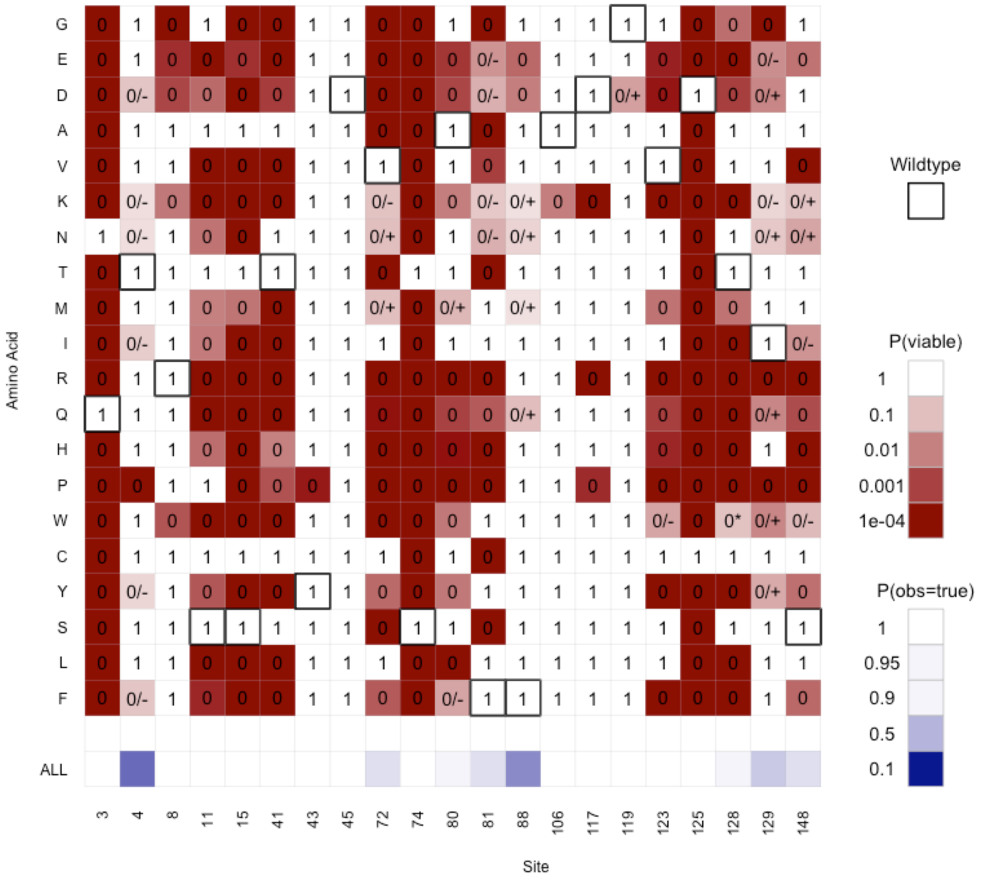


**Supplementary figure S1.** Results of viability analysis identifying 223 viable and 196 inviable genotypes. Red to white heatmap shows the probability that a variant is viable based on NNN plaque sequencing data. 1=observed in NNN mutagenesis, 0= unobserved in NNN mutagenesis. Unobserved variants from NNN mutagenesis with P(viable) > 0.01 were directly made one codon at a time (singles): 0/+ were recovered, and thus viable; 0/- were not recovered and assumed inviable. 0* (at variant 128W) is the one unobserved variant where P(viable) > 0.01 that we failed to attempt to synthesize as a single; the data point was removed. The bottom row (in blue) shows the probability $P_{obs=true}$ before singles were synthesized.

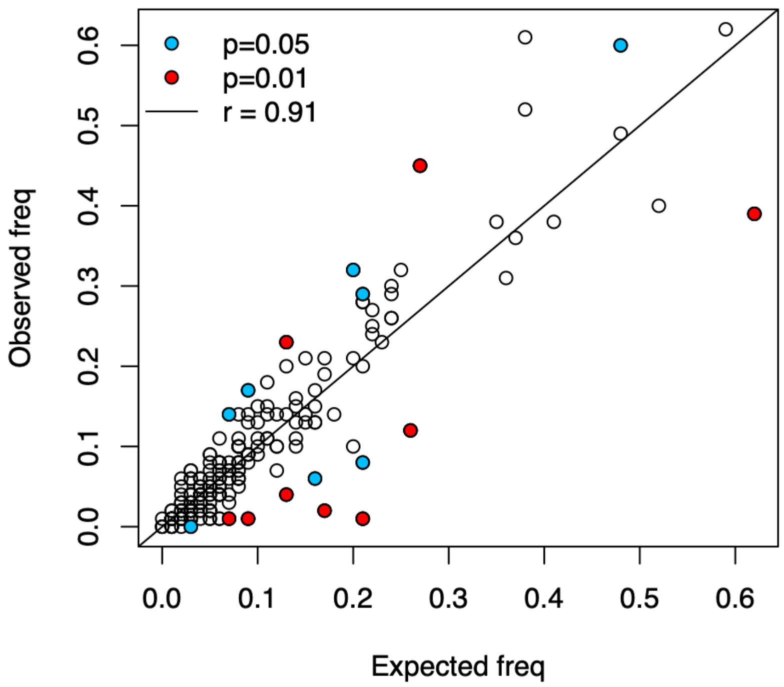


**Supplementary figure S2.** Expected and observed amino acid variant counts across all 21 sites are highly correlated (r = 0.91). Significant deviations are indicated in blue and red. Expected frequency is based on Illumina sequencing of the DNA mutational library. Observed frequency is based on Sanger sequencing of individual plaques.

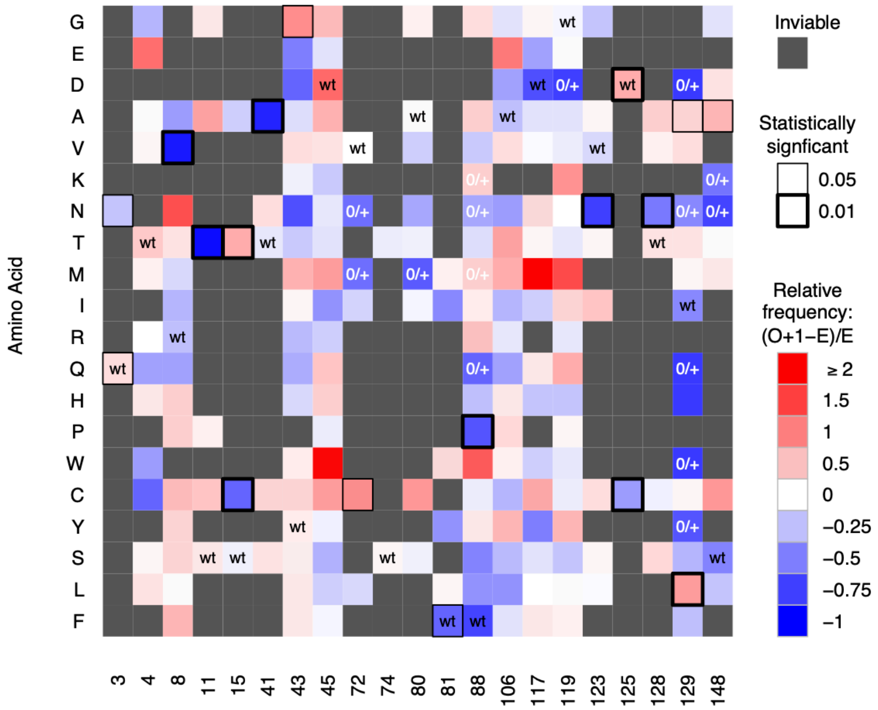


**Supplementary figure S3**. Frequency of variants observation compared to expectation from the ligation mix frequencies. Putatively inviable variants are grayed out. Bold boxes indicate significant deviations from expectation. One observation has been added to each viable variant to moderate small sample size effects. Note, sites 3 and 125 only have two observed amino acids, making it impossible to discern which variant is driving the departure from expectation.

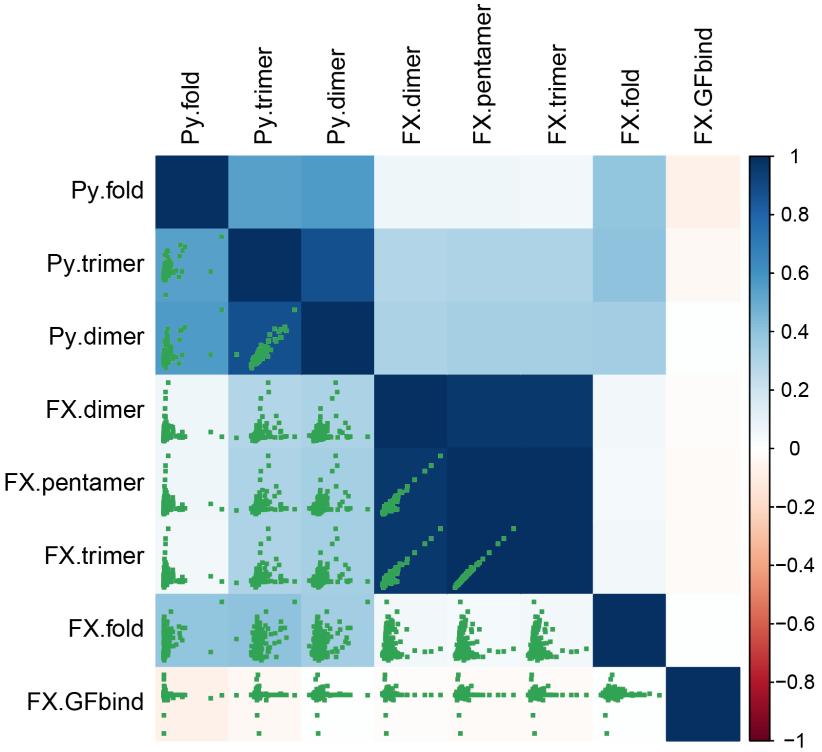


**Supplementary figure S4.** A comparison of the eight modeling methods used in this paper. “Py” and “FX” stand for PyRosetta and FoldX, respectively. Strength of correlation is represented by the color of the squares. Dot plots for only the 21 sites that we mutated are also shown.

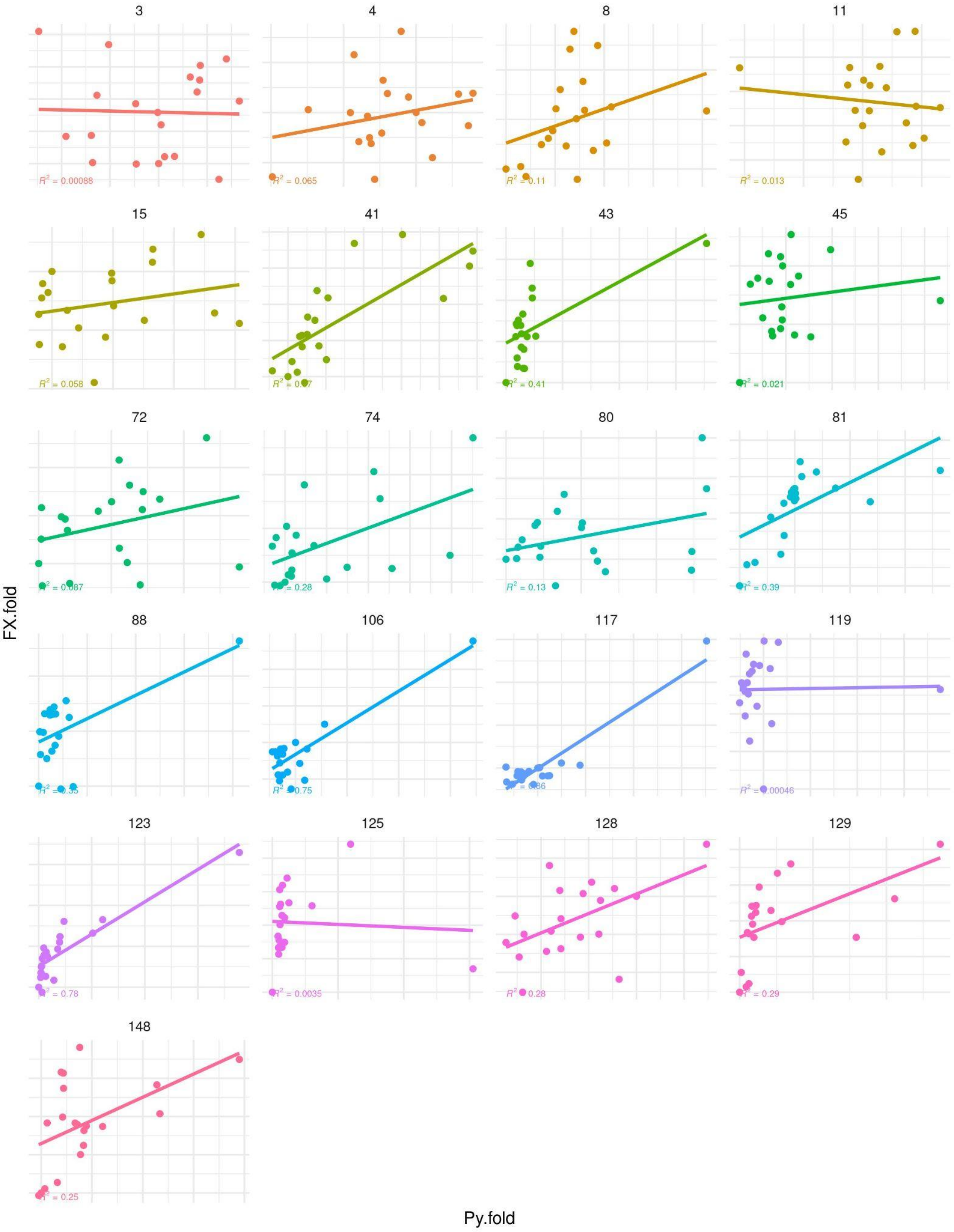


**Supplementary figure S5.** Correlation between FoldX folding and PyRosetta folding ΔΔG values for each residue. Correlation coefficients are shown.

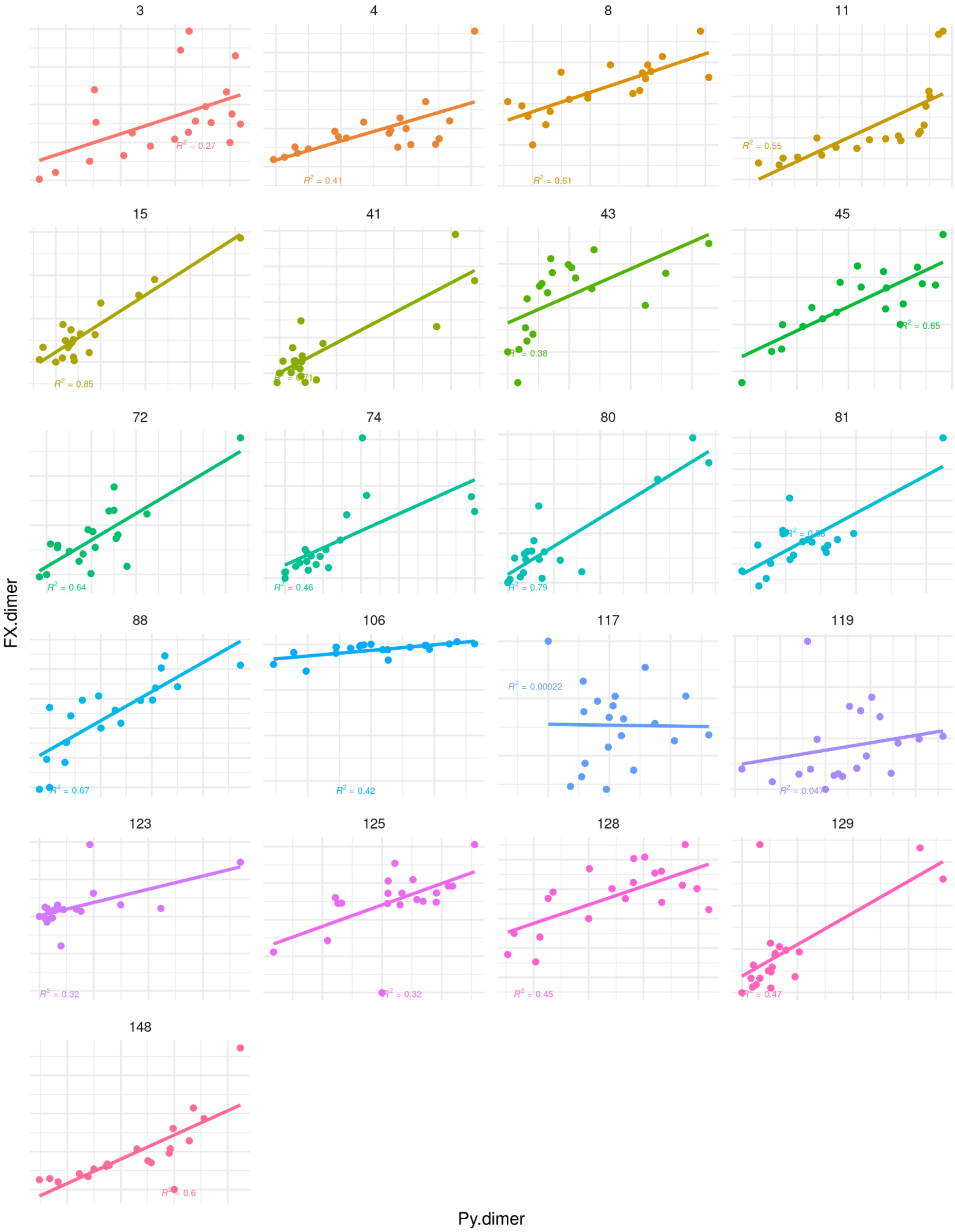


**Supplementary figure S6.** Correlation between FoldX binding and PyRosetta binding ΔΔG values for each residue. Correlation coefficients are shown.

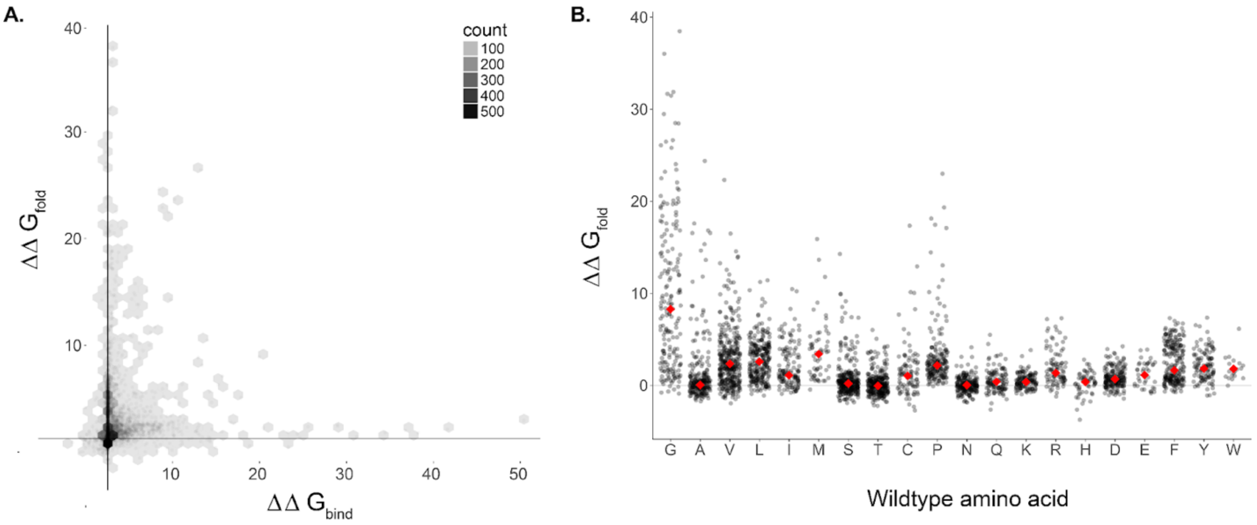


**Supplementary figure S7.** Predicted effects of amino acid substitutions on protein G binding and folding stability (A). Folding effects vary between starting amino acids (B). For example, changing a glycine to anything else is predicted to have large impact on protein folding. Means are shown in red. Calculations from FoldX predictions.

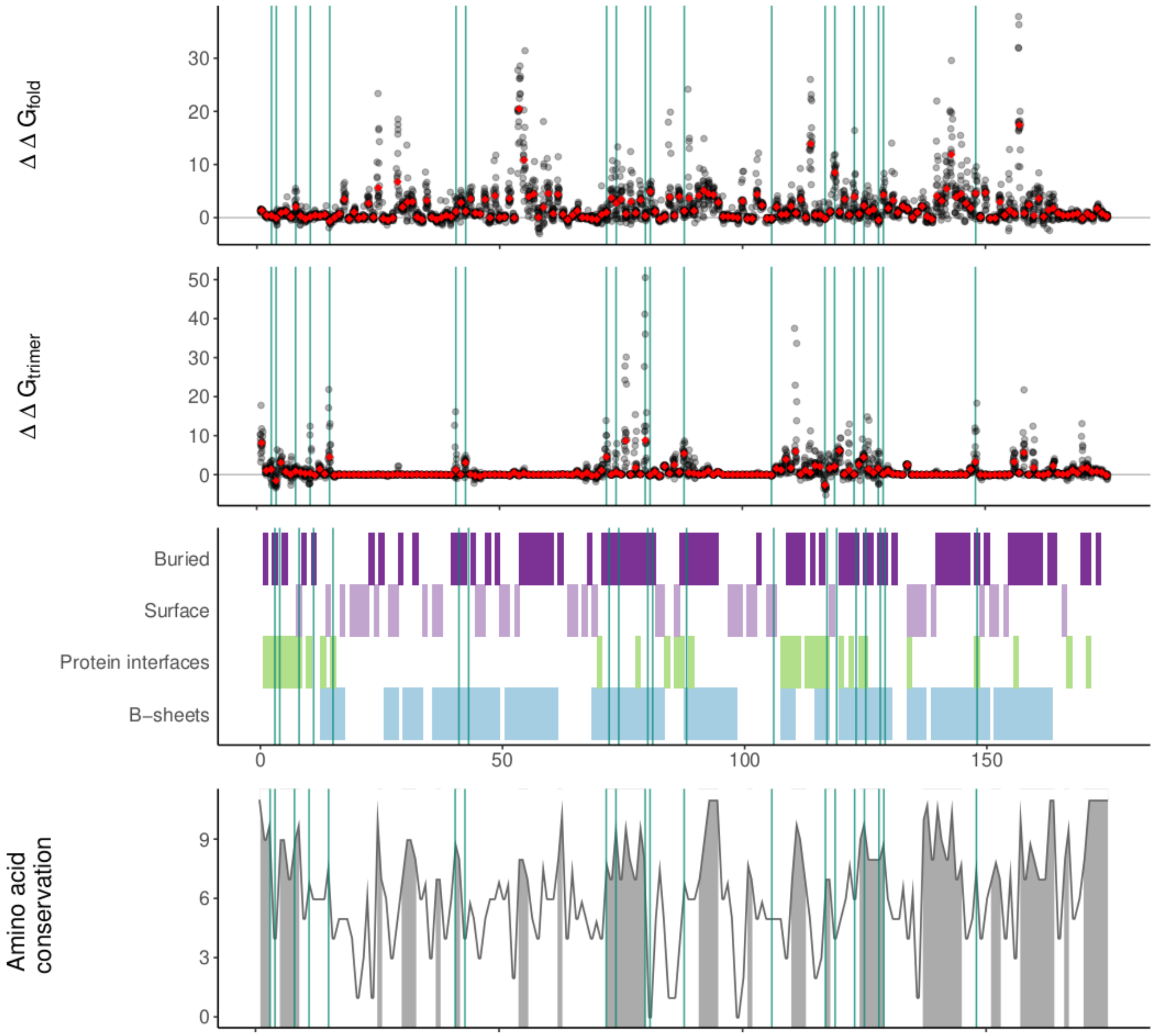


**Supplementary figure S8**. Properties of the ΦX174 G protein. Conservation is calculated according to Livingstone and Barton (1993). The predicted effects of all possible amino acid mutations (black points) on the folding and binding stabilities of G using FoldX. The per-site mean effects is shown with red points. Various structural properties of the G protein are also shown. Residue accessibility was calculated using (http://cib.cf.ocha.ac.jp/bitool/ASA/). Protein interfaces and secondary structure locations were taken from McKenna, Ilag, and Rossman (1994).

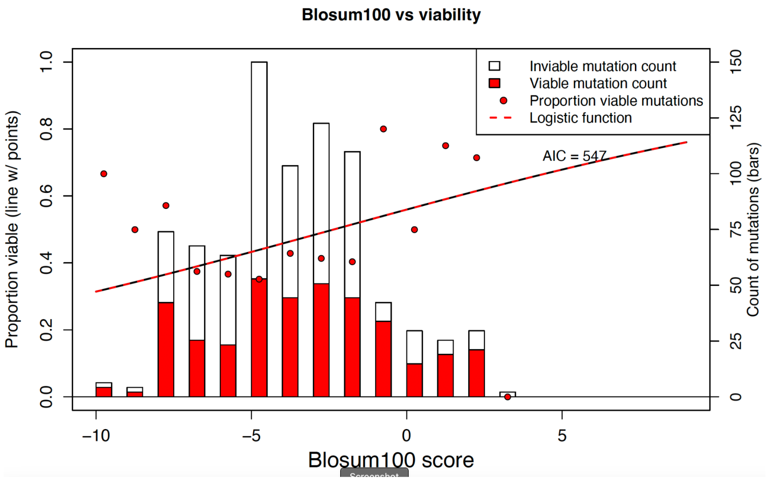


**Supplementary figure S9.** Substitutions with high Blosum100 scores are more likely to be tolerated. Histogram showing the proportion (y-axis) and number (secondary y-axis) of viable substitutions at different Blosum score intervals. We used a modified logistic regression model (see methods for details).

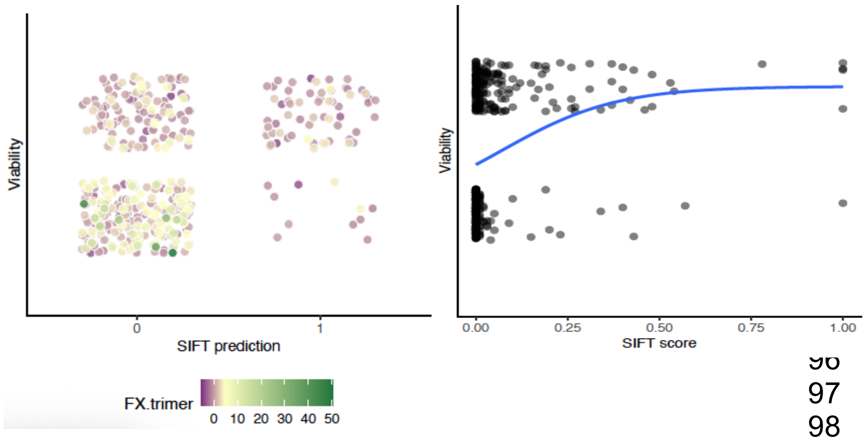


**Supplementary Figure S10**. Correlation between SIFT scores and viability. Left plot bins substitutions into binary categories; viable (y=1) vs. inviable (y=0) and SIFT predicted viable (x=1) vs. SIFT predicted inviable (x=0). The plot on the right shows the distribution of SIFT scores (0-1). We choose the suggested value of 0.05 and above for predicted viable mutants.

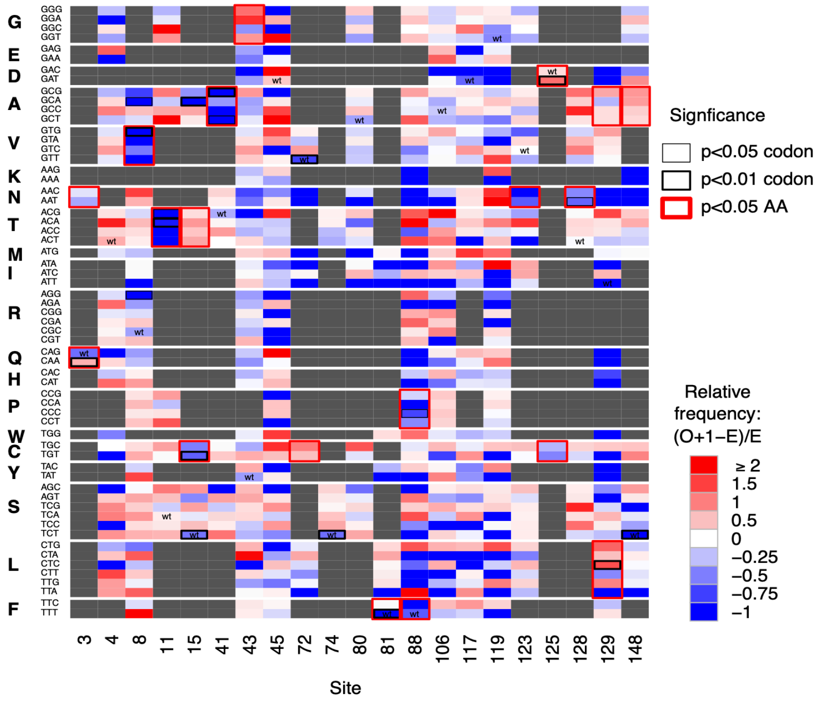


**Supplemental figure S11**. Instances where codon and amino acids are observed in plaque counts more or less than expected based on sequencing of the initial mutational library. Inviable variants are grayed out. Bold black boxes indicate significant codon deviations from expectation. Red boxes indicate significant deviations at the amino acid level. When an amino acid deviates from expectation, the codons also generally do, thus, **Supplementary figure S12** accounts for this by calculating within amino acid codon deviations. One observation has been added to each viable variant to moderate small sample size effects. In general, the figure shows that most of the amino acid deviations are not driven by codon-level effects.

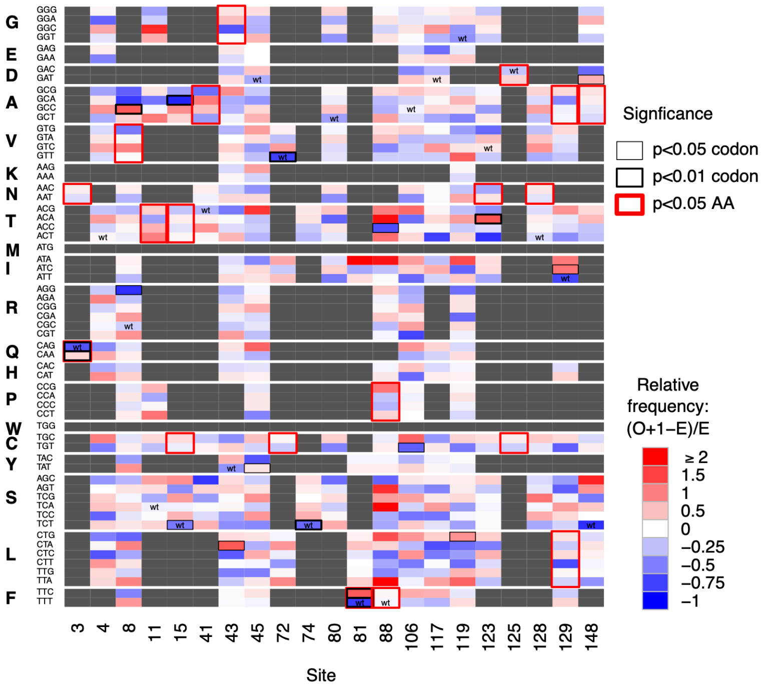


**Supplemental figure 12**. Instances where codon and amino acids are observed in plaque counts more or less than expected based on sequencing of the initial mutational library. Inviable variants are grayed out. Bold black boxes indicate significant deviations from expectation. Red boxes indicate significant deviations at the amino acid level. One observation has been added to each viable variant to moderate small sample size effects. In general, the figure shows that most of the amino acid deviations are not driven by codon-level effects. Significance was determined using parametric bootstrapping with 1000 reps.

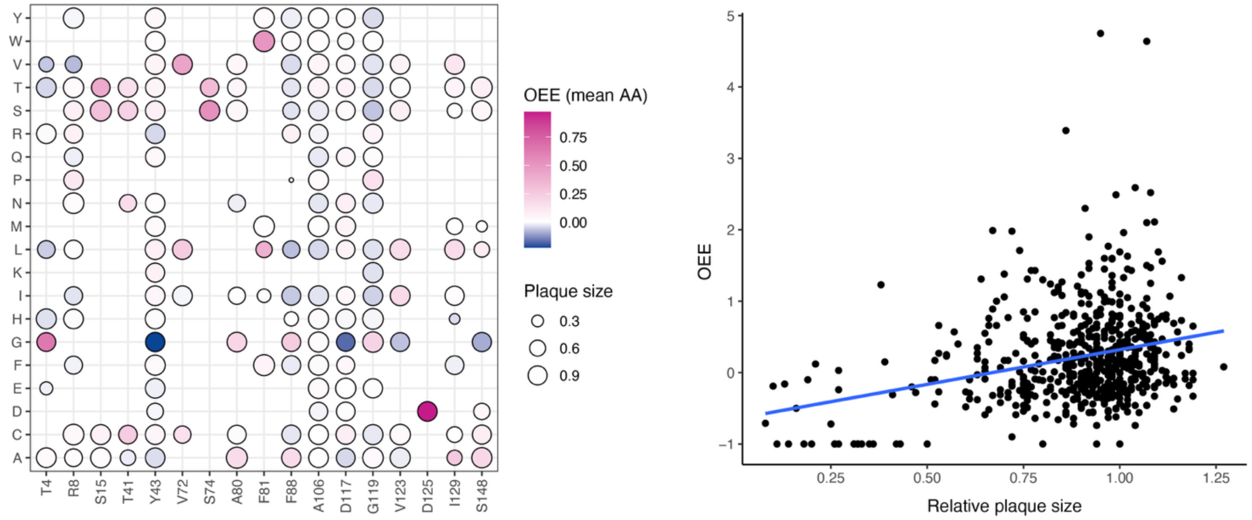


**Supplementary figure S13**. Plaque size variation among mutant genotypes. Left plot shows the size of plaques and the deviation from expected observation frequencies (Observed-Expected / Expected). The average plaque size of all codons was used for the plot on the left (amino acid plot). The plot on the right shows a correlation was observed between plaque size and observation frequency (linear model, p=1.2E-12, $r^2$=0.079). The points along the y=-1 line are variants that were not observed in our plaque picking but were verified to be present in the starting library. A weaker but still significant relationship between OEE and plaque size exists if these variants (linear model, p=6.0E-5, $r^2$=0.027).

**Supplementary table S1.** Fit of different logistic regression models based on AIC and Nagelkerke's pseudo-r2 when residue 119 is included in the dataset. Table X in the main paper shows the same fit statistics when residue 119 is excluded. Models using FoldX (FX) were run with and without predictor error included (except F-G bind for which we had no error estimate); PyRosetta (Py) estimates lack error estimates. While inclusion of error is preferable, we provide fits without as well so fair AIC and $r^2$ comparisons between FX and Py can be made. The best model is FX fold + FX bind: trimer.

| **Model** | **AIC: With (w/o error)** | **$r^2$: With (w/o error)** |
|---|---|---|
| FX F fold | 552 (556) | 0.019 (0.005) |
| FX F bind: trimer | 511 (511) | 0.147 (0.147) |
| FX F bind: dimer | 514 (514) | 0.139 (0.138) |
| FX F bind: pentamer | 519 (515) | 0.123 (0.135) |
| FX F-G bind: pentamers | (558) | (0.000) |
| Py fold | (544) | (0.045) |
| Py bind: trimer | (544) | (0.046) |
| Py bind: dimer | (542) | (0.050) |
| FX fold + FX bind: trimer | 508 (513) | 0.167 (0.152) |
| Py fold + Py bind:trimer | (549) | (0.043) |
